# Hyperactivation of TAK1 causes skeletal muscle pathology reminiscent of inflammatory myopathies

**DOI:** 10.1101/2025.03.21.644671

**Authors:** Meiricris Tomaz da Silva, Anirban Roy, Ashok Kumar

**Author notes:** **Corresponding author**: Ashok Kumar, Ph.D., Institute of Muscle Biology and Cachexia, Department of Pharmacological and Pharmaceutical Sciences Health Building 2, Room 5012, College of Pharmacy, University of Houston Houston, TX 77204-1217.

## Abstract

Loss of skeletal muscle mass and strength is a debilitating consequence of various chronic diseases, inflammatory myopathies, and neuromuscular disorders. Inflammation plays a major role in the perpetuation of myopathy in degenerative muscle diseases. TAK1 is a major signaling protein that mediates the activation of multiple signaling pathways in response to inflammatory cytokines and microbial products. Recent studies have demonstrated that TAK1 is essential for the growth and maintenance of skeletal muscle mass in adult mice. However, the effects of overstimulation of TAK1 activity in the regulation of skeletal muscle mass remain unknown. In the present study, using AAV vectors, we investigated the effect of varying levels of TAK1 activation on skeletal muscle in adult mice. Our results demonstrate that while low levels of TAK1 activation improve skeletal muscle mass, sustained hyperactivation of TAK1 causes myopathy in adult mice. Excessive stimulation of TAK1 manifests pathological features, such as myofiber degeneration and regeneration, cellular infiltration, increased expression of proinflammatory molecules, and interstitial fibrosis. Hyperactivation of TAK1 also upregulates proteolytic systems and various catabolic signaling pathways in skeletal muscle of adult mice. Altogether, our study demonstrates that physiological levels of activation of TAK1 lead to myofiber hypertrophy, whereas its hyperactivation results in myofiber damage and other pathological features resembling inflammatory myopathies.

## Introduction

Skeletal muscle is a highly plastic tissue that undergoes changes in its mass in response to environmental cues. Many anabolic stimuli, such as nutrients and resistance exercise, increase myofiber protein content leading to skeletal muscle hypertrophy. In contrast, catabolic stimuli, such as inactivity, aging, and various chronic diseases augment muscle proteolysis leading to the loss of skeletal muscle mass, commonly known as muscle atrophy or wasting (1–3). The loss of skeletal muscle mass and strength is also a common feature of various neuromuscular disorders, including muscular dystrophy. The skeletal muscle of patients with muscular dystrophy undergoes chronic cycles of myofiber degeneration and regeneration, followed by inflammation, which eventually results in the replacement of muscle tissue with fibrotic tissue (4–6). Similarly, muscle degeneration is also commonly observed in inflammatory myopathies, a group of rare diseases that involve chronic muscle inflammation and weakness (7, 8).

Skeletal muscle mass is governed by a delicate balance between the rate of protein synthesis and degradation, which is regulated by the coordinated activation of multiple signaling pathways (9). The IGF1/Akt/mTOR is a major signaling pathway that augments skeletal muscle mass through the stimulation of protein synthesis and the inhibition of the gene expression of components of the ubiquitin-proteasome system and autophagy (10). By contrast, many proinflammatory cytokines and microbial products cause muscle wasting through the activation of the nuclear factor-kappa B (NF-κB) and p38 MAPK signaling pathways (11, 12). Other cytokines, such as IL-6, mediate muscle wasting through the activation of the STAT3 signaling (13, 14). Furthermore, the TGFβ/Activin/Myostatin subfamily of ligands activates Smad2/3 proteins, which stimulate muscle wasting through augmenting protein degradation (9, 15–19) whereas BMP family ligands activate Smad1/5/8 proteins, which mediate muscle growth and prevents atrophy (16, 18, 20, 21).

TGF-β-activated kinase 1 (TAK1) is an important serine/threonine kinase that mediates the activation of NF-κB, JNK, and p38 MAPK signaling in response to various cytokines, growth factors, and microbial products (22–24). TAK1 forms a signalosome with other proteins, such as TAK1-binding protein (TAB)1 and either TAB2 or TAB3, which bind to the N- and C-terminus of TAK1, respectively (25). Moreover, for its activation, TAK1 undergoes poly-ubiquitination and auto-phosphorylation at the Thr187 residue within its activation domain, followed by phosphorylation at the Thr184 and Ser192 residues (26, 27). TAK1 has been shown to regulate various biological processes, including cell survival, proliferation, differentiation and the inflammatory immune response.

TAK1 plays a crucial role in tissue development, as evidenced by the finding that Tak1-knockout mice exhibit embryonic lethality (28). Studies using tissue-specific Tak1-knockout mice have further highlighted the essential role of TAK1 in the development and homeostasis of various organs (23, 25). Previously, we reported that germline muscle-specific inactivation of TAK1 in mice results in perinatal lethality and tamoxifen-inducible inactivation of TAK1 in the skeletal muscle of adult mice leads to muscle wasting and the development of kyphosis (29). Further evidence supporting the growth-promoting role of TAK1 in skeletal muscle comes from the findings showing that chronic mechanical overload increases TAK1 phosphorylation within its activation domain, along with other positive regulators of muscle mass (29, 30), and that inducible TAK1 inactivation significantly reduces the overload-induced myofiber hypertrophy (29). Notably, forced activation of TAK1 via intramuscular co-injection of TAK1- and TAB1-expressing adeno-associated viruses (AAVs) causes a significant increase in the wet weight and myofiber cross-sectional area in skeletal muscle of adult mice (30). One of the potential mechanisms by which TAK1 induces skeletal muscle growth is through augmenting protein synthesis (29, 30). Intriguingly, a few recent studies have shown that inhibition of TAK1 reduces the amount of fibrosis in dystrophic muscle of mdx (a mouse model of Duchenne muscular dystrophy) mice (31, 32). While the exact mechanisms remain unknown, it is possible that dystrophic muscle presents a chronic injury-associated microenvironment where excessive activation of TAK1 can lead to wasting. There is also a possibility that depending on the level of activation, TAK1 plays distinct roles in the regulation of skeletal muscle mass and fibrosis.

In this study, using AAVs, we have investigated the effects of low and high levels of TAK1 activation on the skeletal muscle in adult mice. Our results demonstrate that while low levels of activation of TAK1 induce myofiber hypertrophy, sustained high levels of activation of TAK1 lead to the loss of skeletal muscle mass, accompanied by inflammation, chronic myofiber degeneration and regeneration, and fibrosis. Furthermore, hyperactivation of TAK1 activates proteolytic systems and multiple catabolic signaling pathways in skeletal muscle of adult mice.

## Results

### Hyperactivation stimulation of TAK1 causes myopathy in adult mice

Previous studies from our group and others have shown that the forced activation of TAK1 requires co-expression of TAK1 and TAB1 protein (30, 33). We have also previously reported that the phosphorylation of TAK1 in plantaris muscle increases about 5-7 folds on day 7 after performing synergistic ablation surgery, a model of chronic mechanical overload (29). Viral vector manufacturing is a challenging process that can lead to inconsistencies in quality and titers and many times virus genome number does not accurately reflects transduction efficiency or amount of transgene expression (34). In this study, we found that intramuscular injection of 2.5 × 10^9^ vg (low dose) AAV6-TAK1/TAB1 leads to about 4-6-fold increase in levels of phosphorylated TAK1 in skeletal muscle. Furthermore, a 10-fold increase in the amounts of AAV6-TAK1/TAB1 (2.5 × 10^10^ vg, high dose) results in ∼40-fold increase in the levels of phosphorylated TAK1 in the skeletal muscle of mice measured after 4 weeks of AAVs injection (**Figure 1A-D**). Furthermore, our analysis showed that there was a significant increase in the wet weight of the TA muscle, but not the GA muscle, in mice expressing low levels of TAK1/TAB1 compared to contralateral GFP-expressing muscle. By contrast, the wet weight of TA or GA muscles injected with high amount of TAK1/TAB1 AAVs was significantly reduced compared to contralateral muscle injected with equivalent dose of AAV6-GFP vector (**Figure 1E, F**). We next generated TA muscle sections and performed H&E staining and morphometric analysis (**Figure 1G**). Consistent with our previously published report (30), proportion of myofiber with larger cross-sectional area (CSA) and average myofiber CSA were significantly higher in TA muscle overexpressing low levels of TAK1/TAB1 compared to contralateral muscle expressing GFP (**Figure 1H, I**). By contrast, TA muscle expressing high levels of TAK1/TAB1 showed signs of severe myopathy, such as centrally nucleated myofibers, cellular infiltrate, and variability in myofiber size (**Figure 1G**; Supplemental **Figure S1**). Moreover, the proportion of myofibers with higher CSA and average myofiber CSA were significantly reduced in TA muscle expressing high levels of TAK1/TAB1 compared to contralateral TA muscle expressing GFP (**Figure 1J, K**).

**FIGURE 1.**
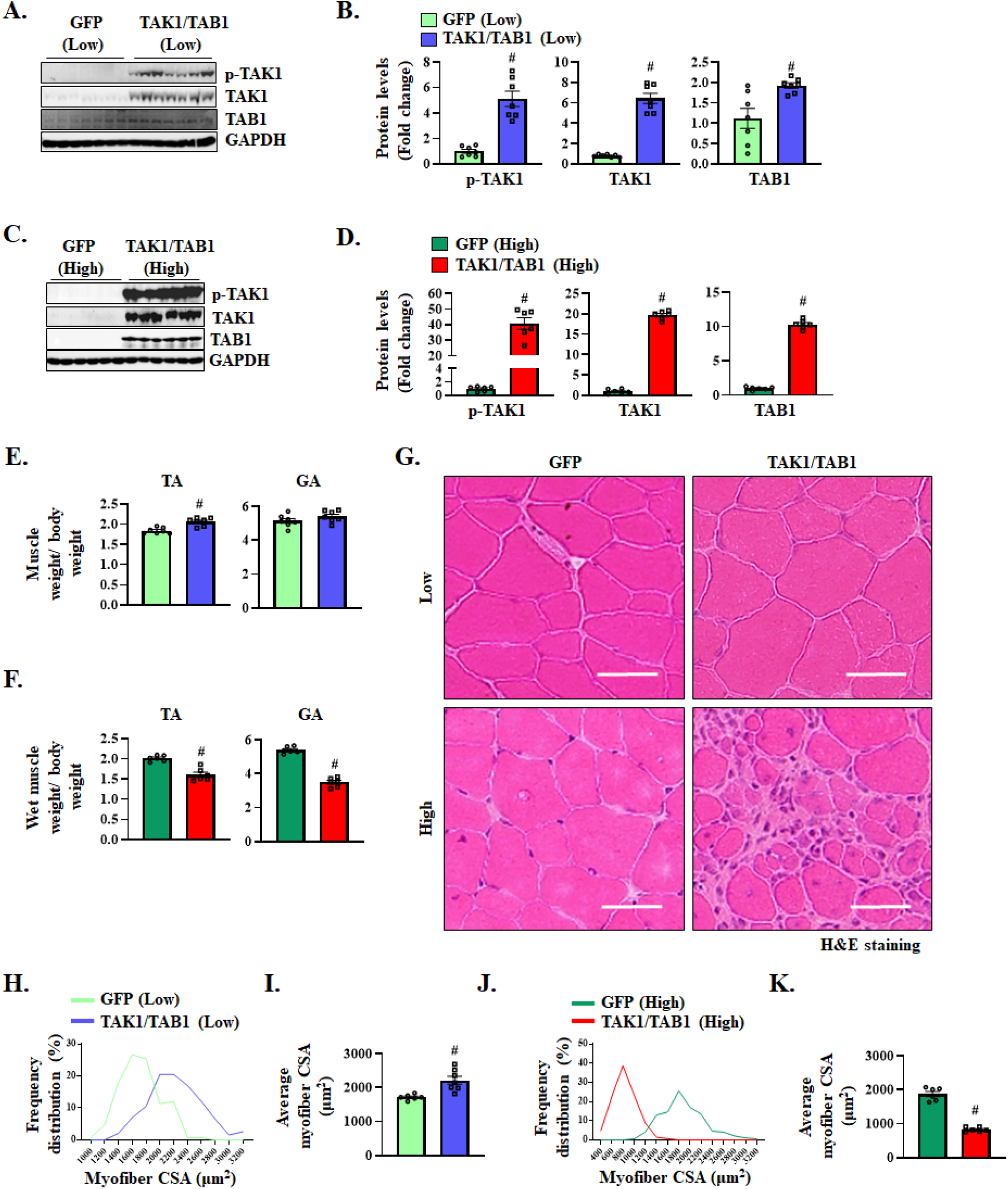
High levels of TAK1 cause myopathy in adult mice. TA or GA muscle of adult wild-type mice was given intramuscular injection of low or high amounts of AAV6-GFP or a combination of AAV6-TAK1 and AAV6-TAB1 and the muscle was analyzed 28 days later. **(A)** Immunoblots, and **(B)** densitometry analysis for protein levels of p-TAK1, TAK1, and TAB1 and unrelated protein GAPDH in GA muscle injected with low amounts of AAV6-GFP or a combination of AAV6-TAK1 and AAV6-TAB1. **(C)** Immunoblots, and **(D)** densitometry analysis for protein levels of p-TAK1, TAK1, TAB1, and GAPDH in GA muscle injected with high levels of AAV6-GFP or AAV6-TAK1/TAB1. TA and GA muscle wet weight normalized by body weight (BW) of mice expressing **(E)** low, or **(F)** high levels of GFP or a combination of TAK1/TAB1. **(G)** Representative photomicrographs of H&E-stained transverse sections of TA muscle of WT mice expressing low or high levels of GFP or a combination of TAK1/TAB1. Scale bar: 50 µm. Quantitative analysis of **(H)** frequency distribution of myofiber cross-sectional area (CSA) and **(I)** average myofiber CSA in TA muscle expressing low levels of GFP or TAK1/TAB1 protein. Quantitative analysis of **(J)** frequency distribution of myofiber CSA and **(K)** average myofiber CSA in TA muscle expressing high levels of GFP or TAK1/TAB1. n= 6-7 mice in each group. All data are presented as mean ± SEM. ^#^p ≤ 0.05, values significantly different from contralateral muscle expressing GFP analyzed by unpaired Student *t* test.

Since high levels of TAK1 cause myopathy similar to inflammatory myopathies or dystrophic muscle, we further analyzed the TA muscle section by performing immunostaining for dystrophin protein. Nuclei were counterstained with DAPI. There was no sign of injury in TA muscle expressing low levels of GFP or TAK1/TAB1 (**Figure 2A**). However, there was drastic increase in the DAPI^+^ nuclei outside the myofibers in the TA muscle expressing high levels of TAK1/TAB1. Furthermore, there were several myofibers filled with mononucleated cells (**Figure 2B**, and Supplemental **Figure S2**), which are typical features of immune cells-mediated myofiber necrosis in inflammatory myopathies (7, 8). Quantitative analysis further showed that there was a significant increase in mid-belly total CSA of whole TA muscle expressing low levels of TAK1/TAB1 and a significant decrease in total CSA of TA muscle expressing high levels of TAK1/TAB1 (**Figure 2C, D**). These results further suggest that while limited activation of TAK1 promotes muscle growth, sustained high levels of TAK1 activity cause skeletal muscle tissue damage.

**FIGURE 2.**
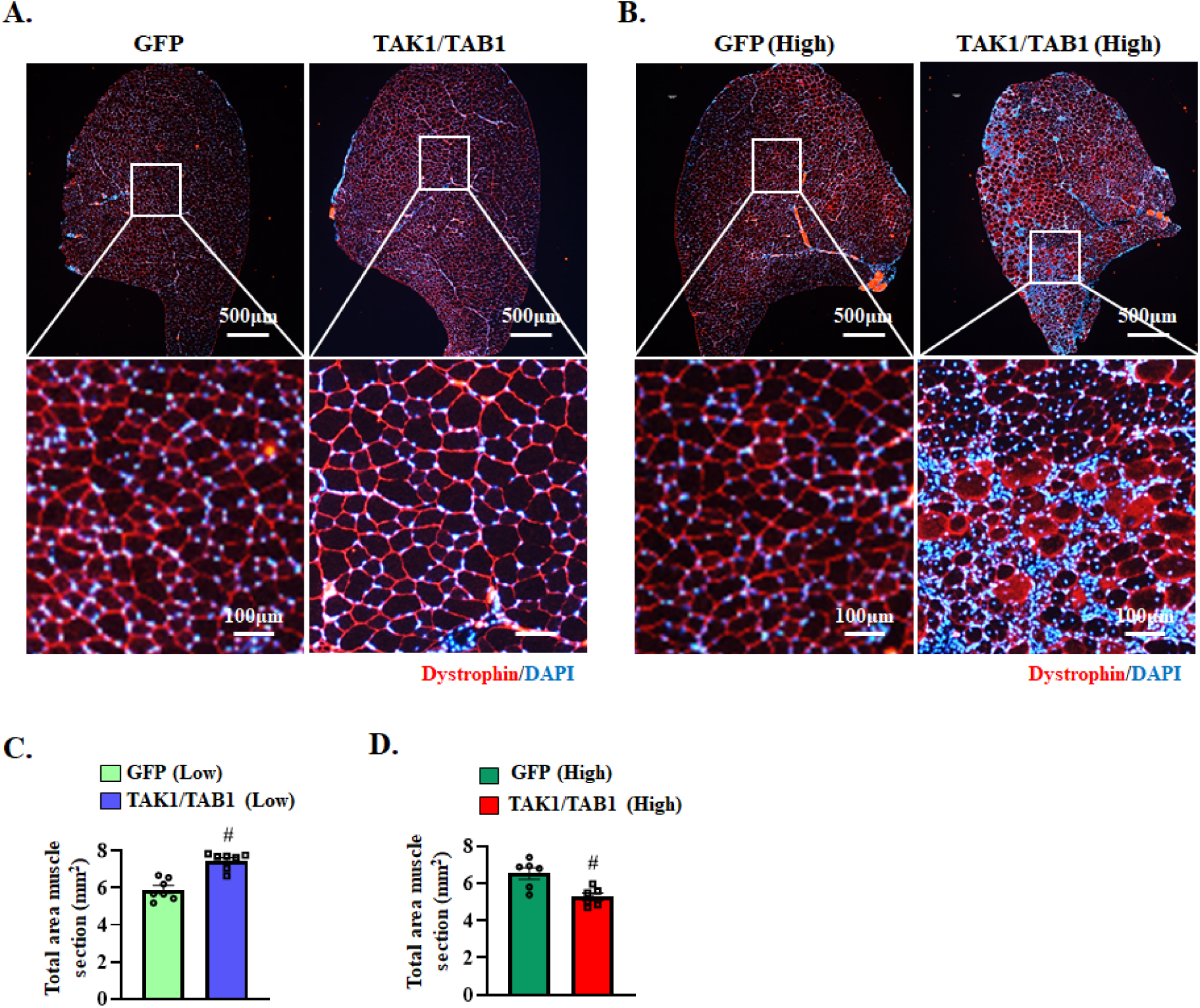
High levels of TAK1 cause myofiber necrosis and inflammation. Representative photomicrographs of transverse sections of TA muscle of mice expressing **(A)** low or **(B)** high levels of GFP or TAK1/TAB1 after immunostaining for dystrophin protein and DAPI staining. Top panel: whole muscle section. Scale bar: 500 µm; bottom panel: magnified view of the selected region. Scale bar: 100 µm. Quantitative analysis of the total area of TA muscle section at mid-belly expressing **(C)** low or **(D)** high levels of GFP or TAK1 and TAB1 protein. n= 6-7 mice in each group. All data are presented as mean ± SEM. #p ≤ 0.05, values significantly different from contralateral muscle expressing GFP analyzed by unpaired Student *t* test.

### High TAK1 activity causes myofiber degeneration and regeneration in skeletal muscle

Skeletal muscle degeneration and regeneration are common features in inflammatory myopathies and in various neuromuscular disorders (4, 35). We next investigated whether high levels of TAK1 activity can induce myofiber damage in skeletal muscle. TA muscle sections of mice injected with low or high titers of TAK1/TAB1 AAVs were immunostained for embryonic myosin heavy chain (eMyHC), a protein that is expressed in newly formed myofibers. The number of eMyHC^+^ myofibers was negligible in TA muscle expressing low levels of GFP, or TAK1/TAB1, or high levels of GFP. By contrast, the eMyHC^+^ myofibers were abundant in TA muscle expressing high levels of TAK1/TAB1 protein (**Figure 3A-D**). We also measured the mRNA and protein levels of some myogenic regulatory factors involved in the regeneration of adult skeletal muscle. There was a significant increase in the mRNA levels of *Myh3* (gene name for eMyHC), *Myod1*, and *Myog* in GA muscle expressing high levels of TAK1/TAB1 compared to contralateral muscle expressing GFP (**Figure 3E**). Furthermore, protein levels of MyoD and Myogenin were significantly increased in GA muscle expressing high levels of TAK1/TAB1 compared to contralateral GFP-expressing GA muscle (**Figure 3F, G**). These results suggest that excessive activation of TAK1 causes muscle injury that stimulates regenerative myogenesis.

**FIGURE 3.**
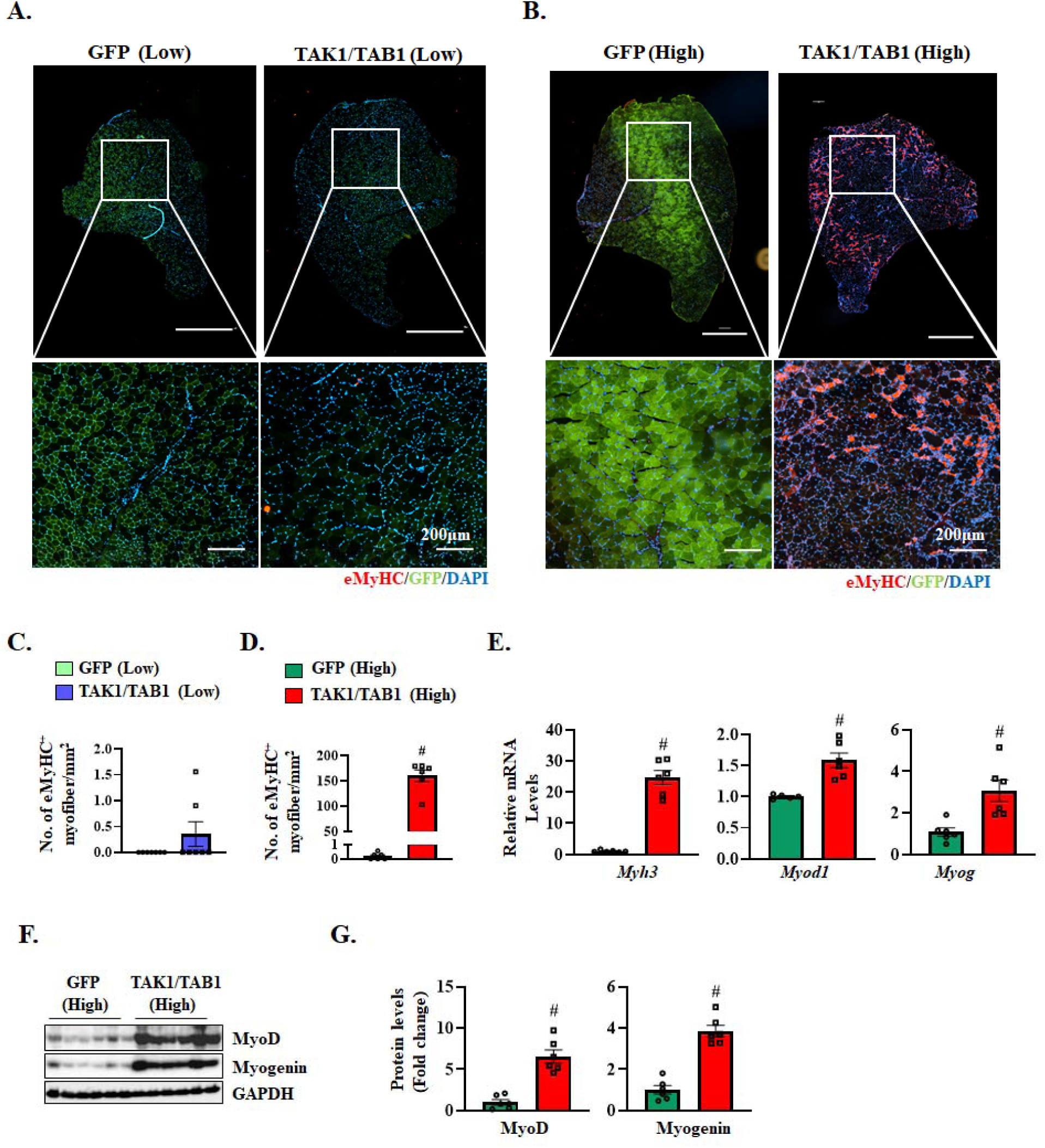
Hyperactivation of TAK1 triggers myofiber degeneration and regeneration in skeletal muscle. Representative photomicrographs of transverse sections of TA muscle of WT mice **(A)** expressing low and **(B)** high levels of GFP or TAK1/TAB1 after immunostaining for eMyHC protein and DAPI staining. GFP in the muscle sections was visualized without any staining. Top panel: whole muscle section; bottom panel: magnified view of the selected region. Scale bar: 200 µm. Number of eMyHC^+^ myofibers in the whole TA muscle section of WT mice expressing **(C)** low and **(D)** high levels of GFP or TAK1/TAB1 protein. **(E)** Relative mRNA levels of *Myh3*, *Myod1* and *Myog* in GA muscle of WT mice expressing high levels of GFP or TAK1/TAB1 protein. **(F)** Immunoblots and **(G)** densitometry analysis of levels of MyoD, Myogenin, and unrelated protein GAPDH in GA muscle of WT mice expressing high levels of GFP or TAK1/TAB1 protein. n= 5-7 mice in each group. All data are presented as mean ± SEM. #p ≤ 0.05, values significantly different from contralateral muscle expressing GFP analyzed by unpaired Student *t* test.

### Hyperactivation of TAK1 activates satellite cells in skeletal muscle of mice

Muscle injury leads to the activation and proliferation of satellite cells, which are essential for myofiber repair (36, 37). Since there was an increase in the number of eMyHC^+^ myofibers in TA muscle of mice expressing high levels of TAK1/TAB1, we next sought to investigate how low or high levels of TAK1 activity affect the number of satellite cells. For this analysis, TA muscle sections were immunostained for Pax7 protein (a marker of satellite cells) and laminin protein (to mark the boundaries of myofibers). Nuclei were counterstained with DAPI. There was no significant difference in the number of satellite cells between TA muscle expressing low levels of GFP or TAK1/TAB1 (**Figure 4A, B**). However, the number of satellite cells significantly increased in TA muscle expressing high levels of TAK1/TAB1 compared to contralateral muscle expressing GFP (**Figure 4A, C**). Quantitative real time-PCR (QRT-PCR) analysis further showed that the mRNA levels of *Pax7* are significantly increased in GA muscle expressing high levels of TAK1/TAB1 compared to contralateral control muscle (**Figure 4D**). Similarly, the protein levels of Pax7 were also significantly elevated in TA muscle with high TAK1/TAB1 activity compared to corresponding control muscle (**Figure 4E, F**). Altogether, these results suggest that sustained high levels of TAK1 causes myofiber degeneration and regeneration that is accompanied by the activation of satellite, cells in skeletal muscle of adult mice.

**FIGURE 4.**
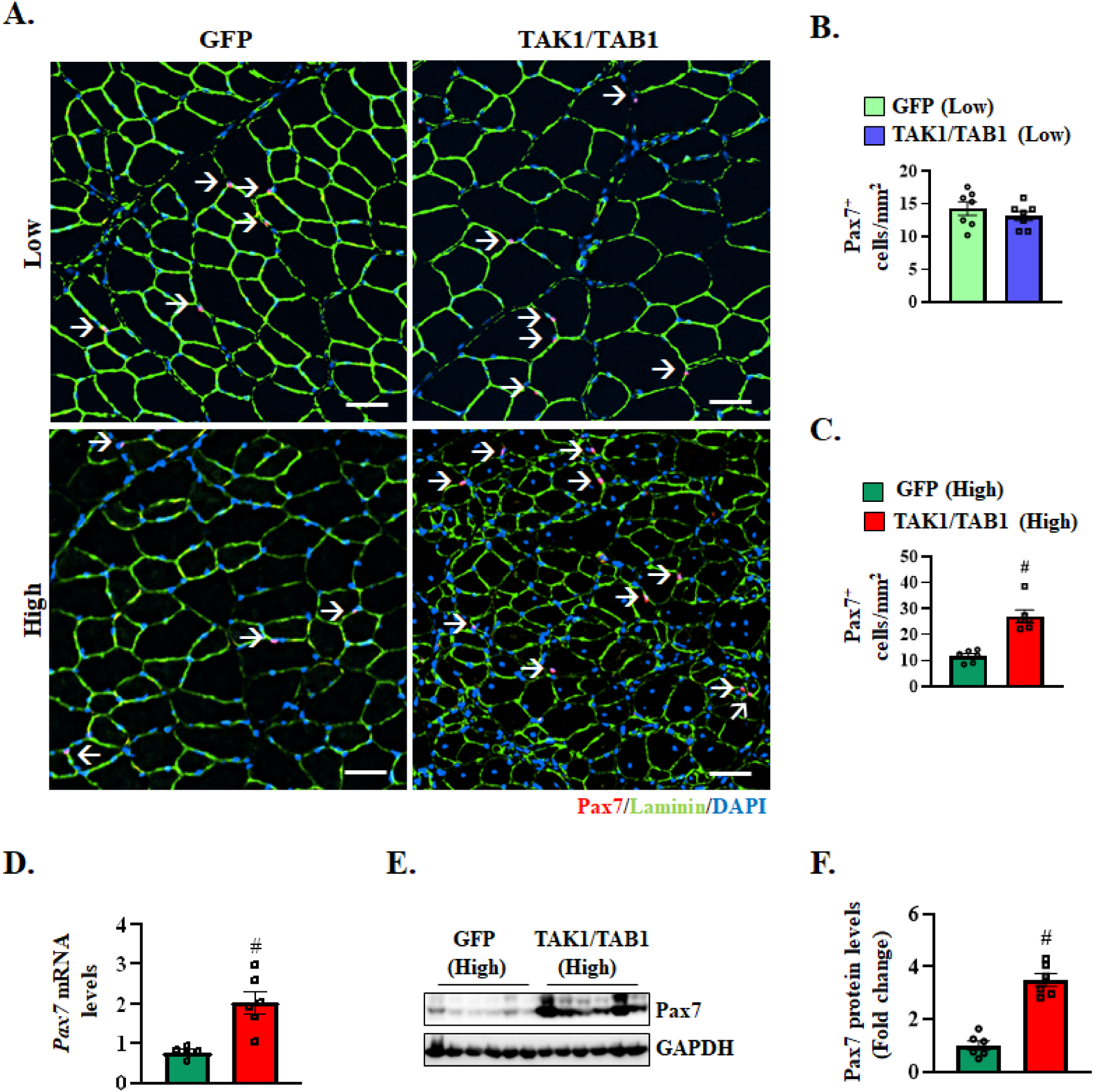
Hyperactivation of TAK1 activates satellite cells in skeletal muscle. **(A)** Representative photomicrograph of transverse sections of TA muscle of WT mice expressing low and high levels of GFP or TAK1/TAB1 protein after immunostaining for Pax7 (red) and laminin (green) protein. Nuclei were identified by staining with DAPI. Quantification of Pax7^+^ cells per unit area in TA muscle section expressing **(B)** low and **(C)** high levels of GFP or TAK1/TAB1 protein. Scale bar: 50 µm. **(D)** Relative mRNA levels of *Pax7* in GA muscle of mice expressing high levels of GFP or TAK1 and TAB1 protein. **(E)** Immunoblots and **(F)** densitometry analysis for protein levels of Pax7 and GAPDH protein in GA muscle of mice expressing high levels of GFP or TAK1/TAB1 protein. n= 5-7 mice in each group. All data are presented as mean ± SEM. #p ≤ 0.05, values significantly different from contralateral injected with AAVV6-GFP analyzed by unpaired Student *t* test.

### Spurious activation of TAK1 causes inflammation and fibrosis in skeletal muscle

TAK1 is an upstream kinase that is known to activate inflammatory signaling pathways in various cell types (22–24). By performing QRT-PCR assay, we examined whether high levels of activation of TAK1 affect the expression of various molecules involved in inflammatory immune response. Results showed that mRNA levels of various cytokines or their receptors, including *Tnfrsf12Aa* (i.e., Fn14), *Tnfrsf1a* (i.e., Tnfr1), *Tnfrsf1b* (i.e., Tnfr2), *Il6* (i.e., IL-6), *Il1b* (i.e., IL-1β), *Tgfb1*, and *Tgfb3* were significantly upregulated in GA muscle expressing high levels of TAK1/TAB1 compared to contralateral muscle expressing GFP. In addition, the gene expression of *Adgre1* (i.e., F4/80), a well-established and widely used marker for murine macrophages, was significantly elevated in GA muscle overexpressing high levels of TAK1/TAB1 compared to GFP-expressing muscle. There was no significant difference in the mRNA levels of *Tnfsf12* (i.e., TWEAK) whereas there was a significant decrease in the mRNA levels of *Tgfb2* in GA muscle expressing high levels of TAK1/TAB1 compared to contralateral muscle overexpressing GFP (**Figure 5A**).

**FIGURE 5.**
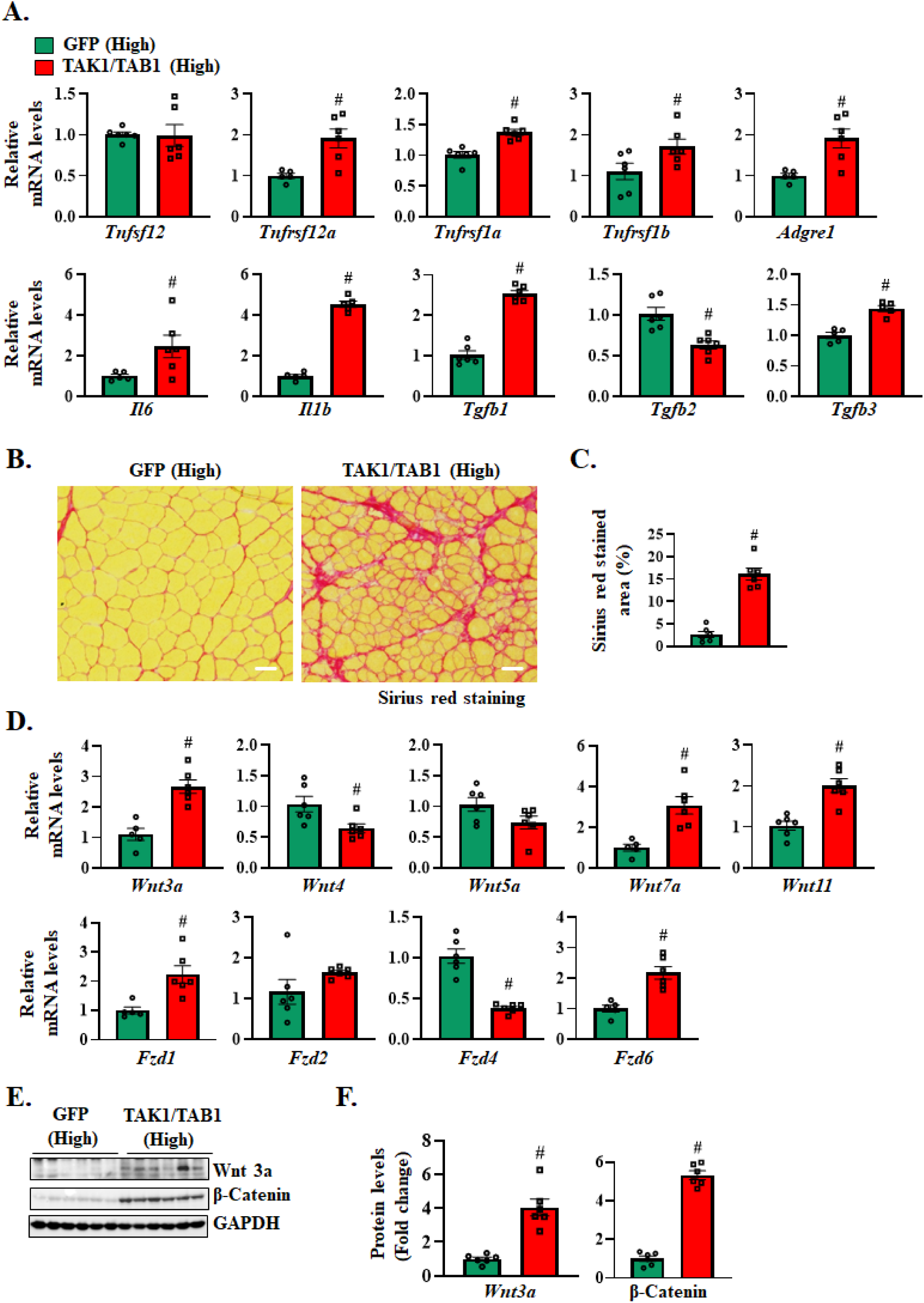
Hyperactivation of TAK1 leads to inflammation and fibrosis in skeletal muscle. **(A)** Relative mRNA levels of *Tnfsf12*, *Tnfrsf12a*, *Tnfrsf1a*, *Tnfrsf1b*, *Adgre1*, *Il6*, *Il1b*,*Tgfb1*, *Tgfb2*, and *Tgfb3* in GA muscle expressing high levels of GFP or TAK1/TAB1. **(B)** Representative photomicrographs of Sirius red-stained TA muscle sections of WT mice expressing high levels of GFP or TAK1/TAB1 protein. Scale bar: 50 μm. **(C)** Quantification of Sirius red-stained area in TA muscle section of mice expressing high levels of GFP or TAK1/TAB1 protein. **(D)** Relative mRNA levels of Wnt ligands: *Wnt3a*, *Wnt4*, *Wnt5a*, *Wnt7a*, and *Wnt11* and Wnt receptors: *Fzd1*, *Fzd2*, *Fzd4*, and *Fzd6* in GA muscle expressing high levels of GFP or TAK1/TAB1 protein. **(E)** Immunoblots and **(F)** densitometry analysis of Wnt3a and β-catenin protein in GA muscle expressing high levels of GFP and TAK1/TAB1 protein. n= 5-6 mice in each group. All data are presented as mean ± SEM. #p ≤ 0.05, values significantly different from contralateral muscle expressing GFP analyzed by unpaired Student *t* test.

Chronic inflammation can lead to the development of tissue fibrosis. TAK1 is also known to contribute to fibrosis in both muscle and non-muscle tissues (31, 32, 38–40). To understand the effect of forced activation of TAK1 on the development of interstitial fibrosis in skeletal muscle, we performed Sirius red staining on TA muscle sections expressing low or high levels of TAK1. Interestingly, a small but significant increase in fibrosis was observed even in TA muscle expressing low levels of TAK1/TAB1 compared to contralateral muscle expressing GFP only (Supplemental **Figure S3**). The amount of fibrosis was drastically increased in TA muscle expressing high levels of TAK1/TAB1 compared to corresponding control muscle (**Figure 5B, C**). Several published studies have demonstrated that the activation of canonical Wnt signaling induces fibrosis in various tissues including skeletal muscle (41–43). Our results demonstrate that the mRNA levels of Wnt ligands (i.e., *Wnt3a*, *Wnt7a*, and *Wnt11*, but not *Wnt4* or *Wnt5a*) and receptors (i.e., *Fzd1* and *Fzd6*, but not *Fzd2* and *Fzd4*) were significantly elevated in GA muscle expressing high levels of TAK1/TAB1 compared to corresponding control muscle (**Figure 5D**). Furthermore, there was also a significant increase in the total levels of Wnt3a and β-catenin, the markers of canonical Wnt signaling, in GA muscle expressing high levels of TAK1/TAB1 compared to contralateral control muscle expressing GFP (**Figure 5E, F**). These results suggest that hyperactivation of TAK1 causes inflammation and fibrosis in skeletal muscle of adult mice.

### Hyperactivation of TAK1 activates proteolytic systems in the skeletal muscle of adult mice

Ubiquitin-proteasome system (UPS) and autophagy are two major proteolytic systems which are responsible for muscle proteolysis in various catabolic conditions (3). We first investigated whether the high levels of TAK1/TAB1 affect the amounts of ubiquitin-conjugated proteins in skeletal muscle of adult mice. Results showed that the mRNA levels of muscle specific E3 ubiquitin ligases, *Fbxo32* (i.e., MAFbx), *Trim63* (i.e., MuRF1), and *Fbxo30* (i.e., MUSA1) were significantly increased in GA muscle expressing high levels of TAK1/TAB1 compared to GFP-expressing control muscle (**Figure 6A**). Moreover, there was also a significant increase in the levels of ubiquitin-conjugated proteins and MuRF1 protein in GA muscle expressing high levels of TAK1/TAB1 compared to contralateral muscle expressing GFP (**Figure 6B, C**). By performing SUnSET assay, we also measured the rate of protein synthesis in GA muscle expressing high levels of TAK1. There was a significant increase in the rate of protein synthesis in GA muscle with hyperactivation of TAK1 compared to corresponding GA muscle expression GFP (**Figure 6B, C**). Since low levels of activation of TAK1 induces myofiber hypertrophy, we also measured the markers of protein synthesis and degradation in GA muscle expressing lower levels of TAK1/TAB1. There was no significant difference in the levels of ubiquitin-conjugated proteins, MAFbx, or MuRF1 between skeletal muscle expressing low levels of GFP or TAK1/TAB1 in GA muscle of mice. However, the levels of puromycin-conjugated peptides were significantly higher in GA muscle expressing low levels of TAK1/TAB1 compared to contralateral control muscle expressing GFP (Supplemental **Figure S4**). These results suggest that low levels of activation of TAK1 promotes protein synthesis without having any effect on the rate of protein degradation whereas hyperactivation of TAK1 augments both protein synthesis and degradation.

**FIGURE 6.**
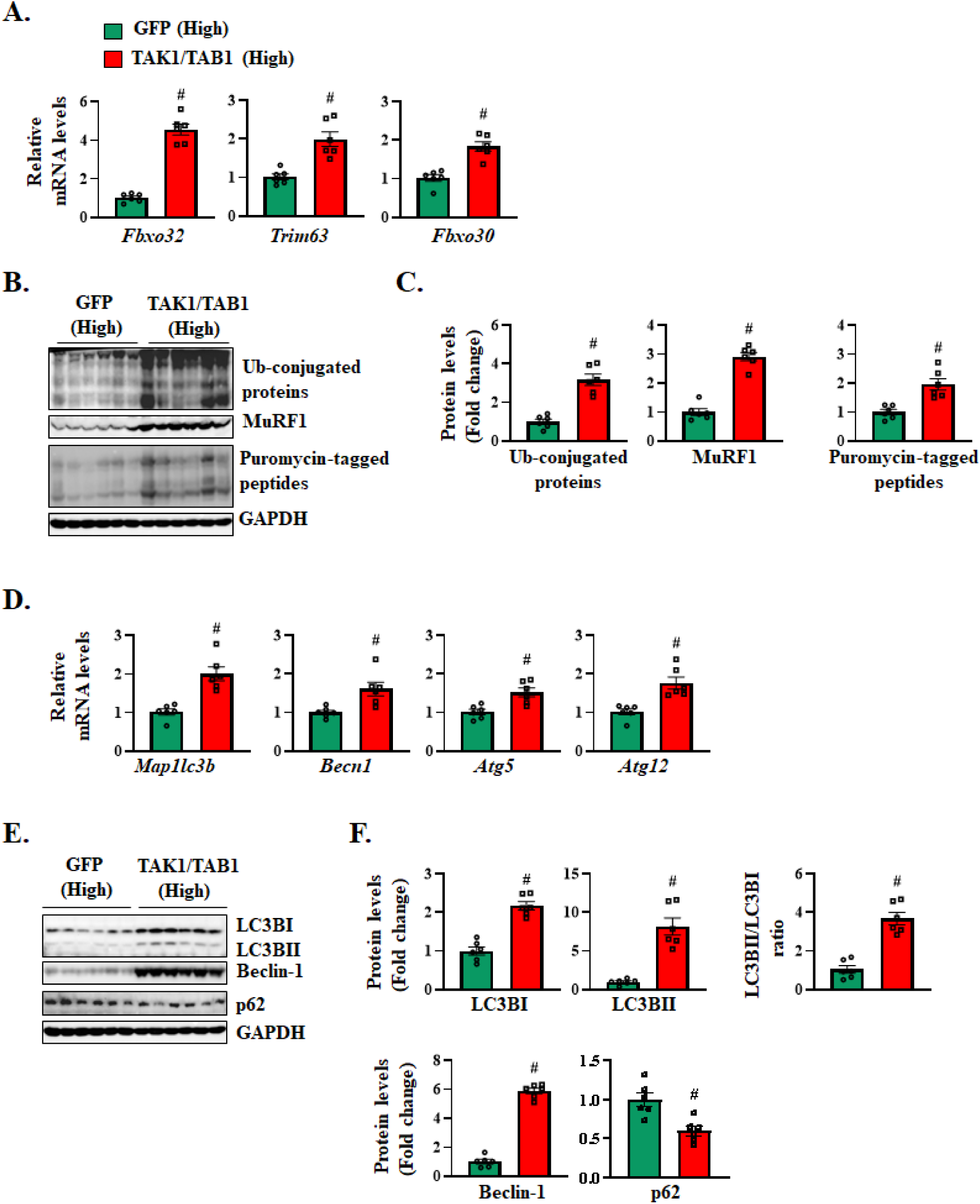
Activation of UPS and autophagy in skeletal muscle with high TAK1 activity. **(A)** Relative mRNA levels of muscle-specific E3 ubiquitin ligases *Fbxo32*, *Trim63*, *Fbxo30* in GA muscle of WT mice expressing high levels of GFP or TAK1 and TAB1 protein. **(B)** Immunoblots and **(C)** densitometry analysis of levels of Ubiquitin (Ub)-conjugated proteins, MuRF1, puromycin-tagged protein, and GAPDH in GA muscle expressing high levels of GFP or a combination of TAK1 and TAB1 protein. **(D)** Relative mRNA levels of autophagy markers *Map1lc3b*, *Becn1*, *Atg5* and *Atg12* in GA muscle expressing high levels of GFP or a combination of TAK1 and TAB1 protein. **(E)** Immunoblots and **(F)** densitometry analysis for protein levels of LC3B I and II, p62, Beclin1, and GAPDH in GA muscle with high levels of GFP or TAK1/TAB1 protein. n= 6 mice in each group. All data are presented as mean ± SEM. #p ≤ 0.05, values significantly different from contralateral muscle expressing GFP analyzed by unpaired Student *t* test. Western blots for B and E were performed contemporaneously.

We next measured the markers of autophagy in GA muscle expressing high levels of GFP or TAK1/TAB1. There was a significant increase in the mRNA levels of *Map1lc3b* (i.e., LCB3B), *Becn1* (i.e., Beclin1), *Atg5*, and *Atg12* in GA muscle with high TAK1 activity compared to control muscle (**Figure 6D**). During autophagy, LC3-I protein undergoes lipidation resulting in the formation of LC3-II, which is then recruited to the membranes of nascent autophagosomes (44). Western blot analysis showed that there was a significant increase in the protein levels of both LC3BI and II and ratio of LC3BII/I in GA muscle overexpressing high levels of TAK/TAB1 compared to contralateral muscle expressing GFP. Furthermore, the protein levels of Beclin-1 were highly elevated in GA muscle with high TAK1 activity. Since p62 is itself an autophagy substrate, it is used as a marker for autophagic flux, along with LC3B (44). Consistent with the increase in LC3BII/I ratio, we found that levels of p62 were significantly reduced in GA muscle overexpressing TAK1/TAB1 compared to control muscle (**Figure 6E, F**). Altogether, these results suggest that hyperactivation of TAK1 stimulates UPS and autophagy in skeletal muscle of adult mice.

### Hyperactivation of TAK1 stimulates the activity of catabolic signaling pathways in skeletal muscle

Multiple signaling pathways regulate skeletal muscle mass in different catabolic conditions (3, 9). By performing western blots, we measured the phosphorylation and total levels of key components of different signaling pathways implicated in skeletal muscle wasting in various conditions. TAK1 is well known to mediate the activation of NF-κB, JNK1/2, and p38 MAPK signaling in response to various proinflammatory cytokines and microbial products. Our analysis showed that the levels of phosphorylated and total IκBα and p65 (widely used markers of canonical NF-κB signaling) and total levels of p100 and p52 protein (markers of non-canonical NF-κB signaling) were significantly elevated in GA muscle overexpressing high levels of TAK1/TAB1 compared to contralateral muscle expressing GFP (**Figure 7A, B**). Furthermore, the levels of phosphorylated and total JNK1/2 and p38 MAPK were also significantly increased in GA muscle overexpressing TAK/TAB1 compared to controls (**Figure 7A, B**).

**FIGURE 7.**
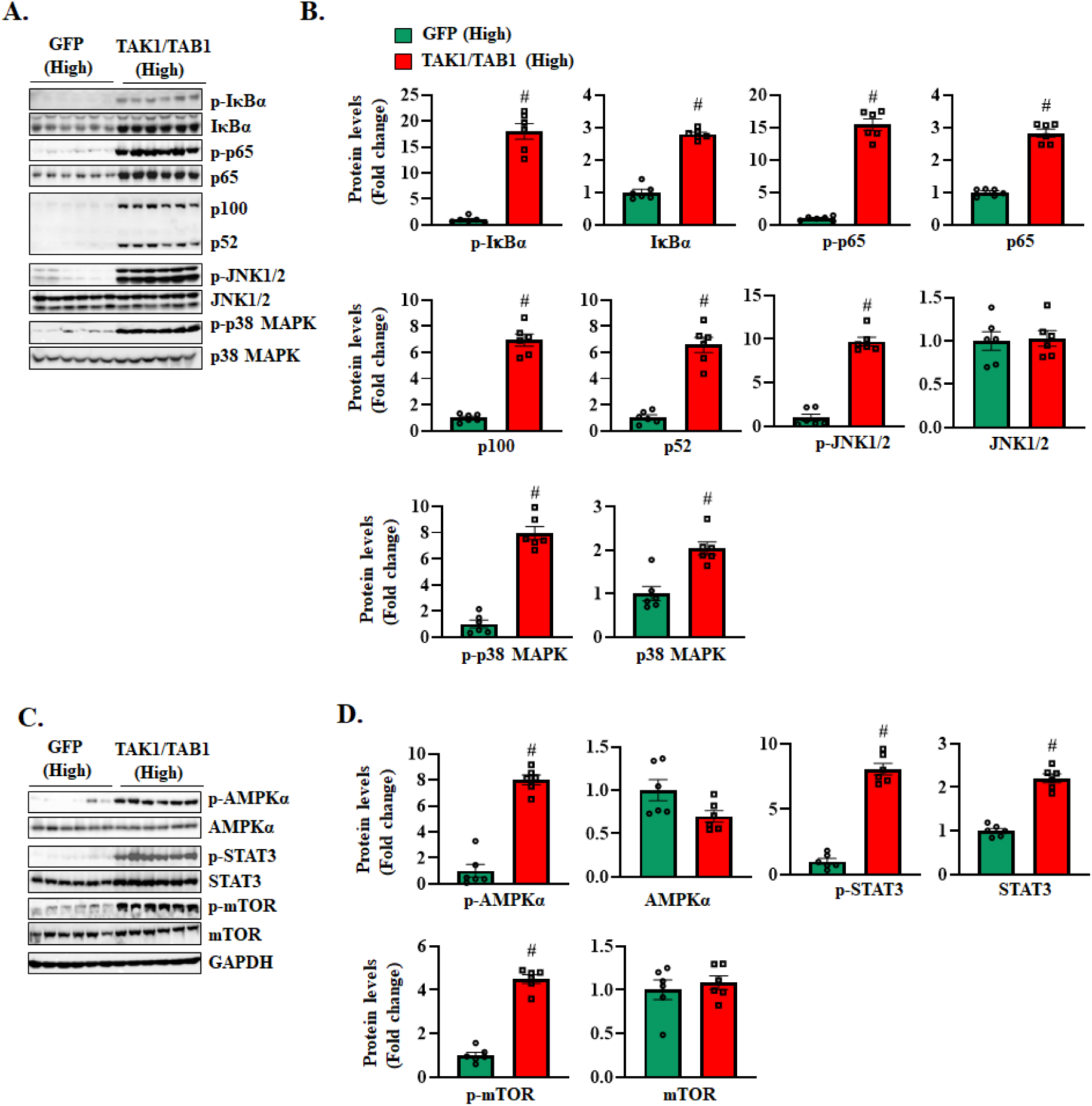
High TAK1 activity stimulates catabolic signaling in adult skeletal muscle. **(A)** Immunoblots and **(B)** densitometry analysis for protein levels of p-IκBα, IκBα, p-p65, p65, p100-p52, p-JNK, JNK, p-p38, and p38 in GA muscle expressing high levels of GFP or TAK1/TAB1 protein. **(C)** Immunoblots and **(D)** densitometry analysis of p-AMPKα, AMPKα, p-STAT3, STAT3, p-mTOR, and mTOR protein in GA muscle expressing high levels of GFP or TAK1/TAB1 protein. n= 6 mice in each group. All data are presented as mean ± SEM. #p ≤ 0.05, values significantly different from contralateral muscle expressing GFP analyzed by unpaired Student *t* test.

Activation of AMPK exacerbates muscle wasting through induction of autophagy and inhibition of mTOR-dependent protein synthesis (45–49). Furthermore, in many conditions, including cancer-induced cachexia, IL-6 induces skeletal muscle wasting through the activation of STAT3 signaling (13, 14). Recent studies have also demonstrated that chronic activation of mTOR causes muscle atrophy and myopathy in many conditions (50–52). Western blot analysis showed that the levels of phosphorylated AMPK and mTOR and phosphorylated and total STAT3 protein were also significantly upregulated in GA muscle with high levels of TAK1 activity (**Figure 7C, D**). Taken together, these results suggest that hyperactivation of TAK1 results in the activation of multiple catabolic signaling pathways in skeletal muscle of adult mice.

### Hyperactivation of TAK1 disrupts Smad signaling in skeletal muscle

Skeletal muscle mass is regulated by coordinated activation of Smad2/3 and Smad1/5/8 signaling pathways. TGFβ family members, including GDF8, GDF11, and Activin A induce muscle wasting through the activation of Smad2/3 signaling. In contrast, BMP family ligands induce muscle growth through the activation of Smad1/5/8 signaling. We investigated whether forced activation of high levels of TAK1 affects the phosphorylation of Smad2/3 or Smad1/5/8 in skeletal muscle of adult mice. Interestingly, the levels of phosphorylated and total Smad2 and Smad1/5/8 were significantly increased in GA muscle expressing high levels of TAK1/TAB1 compared to contralateral muscle expressing GFP (**Figure 8A, B**). Furthermore, our QRT-PCR analysis showed that gene expression of multiple ligands and receptors of TGFβ/Myostatin/Activin A and BMP signaling pathways was disrupted in GA muscle expressing high levels of TAK1/TAB1 (**Figure 8C**). While the physiological significance remains unknown, these results suggest that hyperactivation of TAK1 deregulates both Smad2/3 and Smad1/5/8 signaling in skeletal muscle of mice.

**FIGURE 8.**
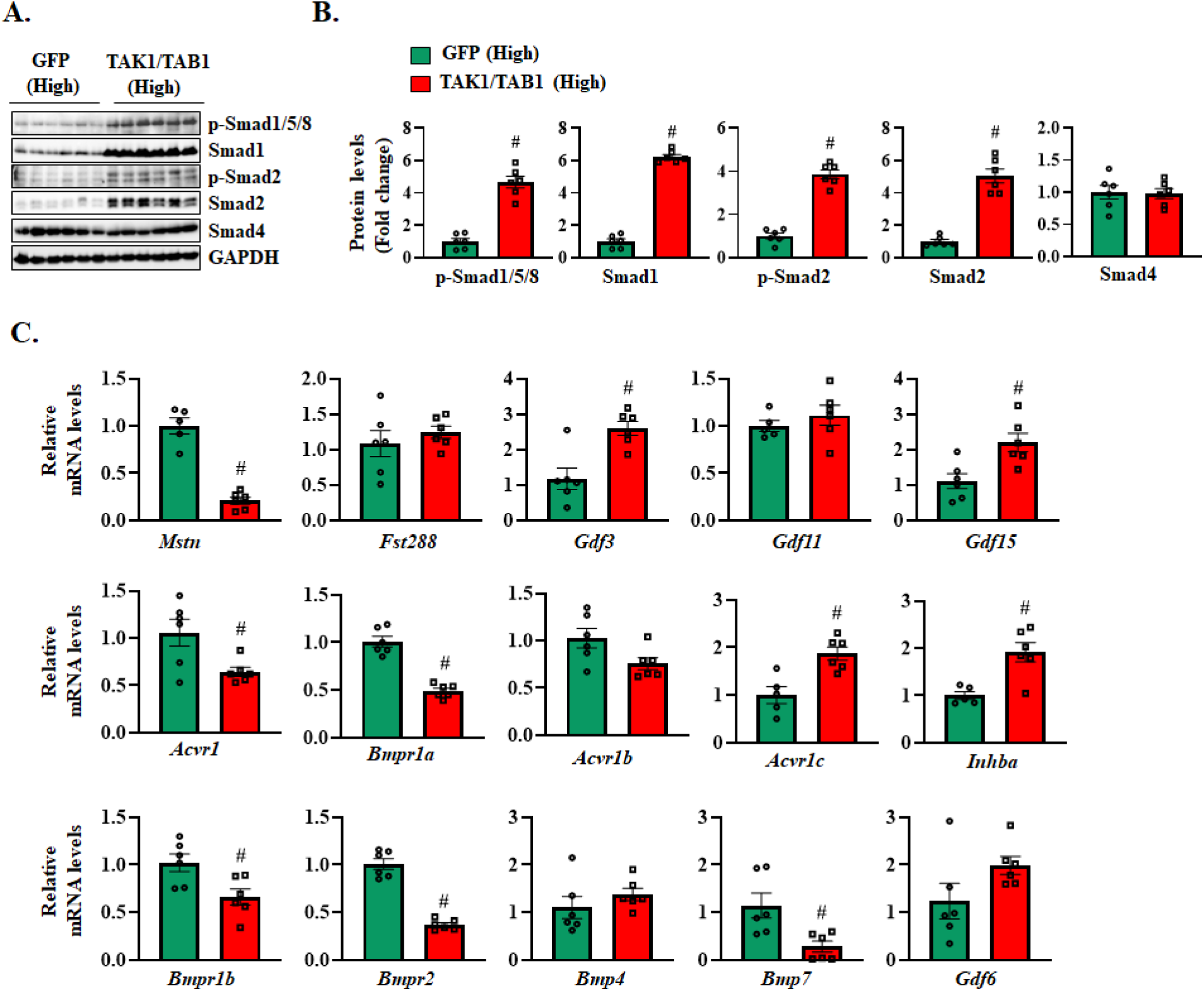
Disruption of Smad signaling in skeletal muscle expressing high levels of TAK1/TAB1 protein. **(A)** Immunoblots, and **(B)** densitometry analysis for protein levels of p-Smad1/5/8, Smad1, p-Smad2, Smad2, and Smad4 in GA muscle expressing high levels of GFP or a combination of TAK1 and TAB1. (C) Relative mRNA levels of *Mstn*, *Fst288*, *Gdf3*, *Gdf11*, *Gdf15*, *Acrv1*, *Bmpr1a*, *Acvr1b*, *Acvr1c*, *Inhba*, *Bmpr1b*, *Bmpr2*, *Bmp4*, *Bmp7*, *Gdf6* in GA muscle of mice expressing high levels of GFP or TAK1/TAB1 protein. n= 5-6 mice in each group. All data are presented as mean ± SEM. #p ≤ 0.05, values significantly different from contralateral muscle expressing GFP analyzed by unpaired Student *t* test.

## Discussion

TAK1 is a major signalosome that regulates different cellular responses through the activation of specific signaling pathways (22–24). Accumulating evidence suggests that TAK1 regulates various aspects of skeletal muscle biology. TAK1 is essential for the maintenance of satellite cell pool and regeneration of skeletal muscle in adult mice (52). TAK1 is also required for post-natal growth and maintenance of skeletal muscle mass in adult animals (53). Moreover, TAK1 promotes myofiber hypertrophy in response to functional overload and its forced activation at low levels causes skeletal muscle growth in adult mice (29, 30). In contrast, there are also several published reports implicating the role of TAK1 in causing inflammation and fibrosis in various tissues, including skeletal muscle (22, 24, 52). We have recently reported that targeted deletion of TAK1 ameliorates muscle histopathology in young mdx mice (54). Other studies have demonstrated that pharmacological inhibition of TAK1 reduces fibrosis in dystrophic muscle of mdx mice (31, 32). However, the role and the mechanisms of action of TAK1 in skeletal muscle physiology and pathophysiology remain incompletely understood.

In this study, we examined the effects of forced low and high levels of TAK1 activation in skeletal muscle of adult mice. Our results demonstrate that while limited activation of TAK1 promotes myofiber growth, hyperactivation of TAK1 results in skeletal muscle pathogenesis in adult mice (**Figure 1**). Overstimulation of TAK1 in skeletal muscle results in muscle degeneration and regeneration, inflammation, and fibrosis, which are commonly observed in inflammatory myopathies (**Figures 1, 2,** and **5**). TAK1 is a major signaling intermediate in many proinflammatory signaling pathways (22). We have found that hyperactivation of TAK1 stimulates the activation of proinflammatory signaling pathways, such as NF-κB and MAPKs and accumulation of inflammatory immune cells in skeletal muscle tissues (**Figures 2** and **Figure 7**). In many pathological conditions, such as inflammatory myopathies, activated immune cells, including T cells and macrophages, infiltrate muscle tissue, leading to myofiber injury through release of inflammatory cytokines and direct cell-mediated cytotoxicity (7, 35, 55). Our results show that the skeletal muscle expressing high levels of TAK1 demonstrates signs of myofiber degeneration and regeneration, which is also evidenced by the presence of eMyHC^+^ myofibers and increased number of satellite cells (**Figure 3** and **Figure 4**). Therefore, it is plausible that the hyperactivation of TAK1 in skeletal muscle increases the levels of various cytokines and chemokines resulting in the recruitment of immune cells, which cause muscle injury through direct or indirect mechanisms.

Our study also suggests that hyperactivation of TAK1 induces myofiber atrophy in skeletal muscle of adult mice. This is evidenced by the findings that the levels of ubiquitinated proteins and the gene expression of muscle-specific E3 ligases, such as MuRF1, MAFbx, and MUSA1 were upregulated in skeletal muscle overexpressing high levels of TAK1/TAB1. In addition, the mRNA and protein levels of markers of autophagy were highly increased in skeletal muscle with high levels of TAK1 activity (**Figure 6**). Since both UPS and autophagy play a major role in muscle proteolysis, their increased activation could be one of the reasons for myofiber atrophy in skeletal muscle expressing high levels of TAK1/TAB1.

Our experiments also demonstrate that hyperactivation of TAK1 disrupts the activation of multiple signaling pathways. NF-kB is a well-known transcription factor that mediates inflammatory immune response through enhancing the gene expression of various proinflammatory cytokines and chemokines (11, 56). Furthermore, proinflammatory cytokines, such as TNFα, TWEAK, and IL-1β and tumor-derived factors induce muscle wasting through the activation of NF-κB (11, 57, 58). Indeed, NF-κB also increases the gene expression of a few components of UPS (11, 59). It has been previously reported that transgenic overexpression of IKKβ, an upstream activator of canonical NF-kB signaling, induces muscle wasting in adult animals (60). Our results demonstrate that the activation of both canonical and non-canonical

NF-κB signaling is significantly increased in skeletal muscle with high levels of TAK1 activity (**Figure 7**). The increased activation of NF-κB signaling could be an important mechanism not only for myofiber atrophy but also for inflammatory immune response observed in skeletal muscle of adult mice expressing high levels of TAK1/TAB1 proteins.

TAK1 also activates JNK and p38 MAPK in mammalian cells (22, 23, 28). While the role of JNK in the regulation of skeletal muscle mass is not clearly understood, there are many reports demonstrating that p38 MAPK mediates muscle atrophy in response to microbial products and tumor-derived factors (12, 61). Our results demonstrate that hyperactivation of TAK1 also increases the phosphorylation of JNK1/2 and p38MAPK in skeletal muscle of adult mice (**Figure 7**). The JAK-STAT3 signaling also stimulates muscle proteolysis in various conditions, including in response to IL-6 cytokine. Similarly, the activation of AMPK is known to contribute muscle atrophy through distinct mechanisms (3, 13, 14, 62). Our results demonstrate increased activation of STAT3 and AMPK in skeletal muscle of mice expressing high levels of TAK1 and TAB1 protein, which could be additional mechanisms for the myofiber atrophy (**Figure 7**). The mechanisms by which TAK1 activates STAT3 or AMPK signaling in skeletal muscle remain completely unknown. However, it is possible that TAK1-mediated signaling crosstalk with the components of JAK-STAT or AMPK pathways. Alternatively, it is possible that the increased expression of specific molecules, such as IL-6 stimulates the activation of these pathways in skeletal muscle. Indeed, the gene expression of IL-6 is significantly upregulated in skeletal muscle with high TAK1 activity (**Figure 5A**).

While mTORC1 was initially considered as a major mechanism of protein synthesis and muscle growth, recent studies have demonstrated that constitutive activation of mTORC1 causes skeletal muscle atrophy and myopathy (50–52). For example, the genetic ablation of the mTORC1 inhibitor TSC1 (tuberous sclerosis complex 1) exacerbates denervation-induced atrophy in mice (63). Moreover, sustained activation of mTORC1 inhibits autophagy in skeletal muscle leading to the onset of myopathy at later stages (64). Similarly, acute activation or suppression of mTORC1 dysregulates autophagy and impairs skeletal muscle homeostasis following denervation (65). Hyperactivation of mTORC1 also contributes to age-related muscle atrophy by enhancing the phosphorylation of STAT3 and increasing the expression of GDF-15 (51). Interesting, we found that the levels of phosphorylated mTOR are significantly increased in skeletal muscle overexpressing high levels of TAK1/TAB1 (**Figure 7**). These elevated levels of mTOR could be another mechanism for the activation of STAT3 and myopathy observed in skeletal muscle of adult mice with chronic high levels of TAK1 activity.

TAK1 is known to activate diverse non-Smad and Smad signaling pathways in response to stimulation by TGFβ or BMP family ligands (52). An earlier study had demonstrated that enzymatically active TAK1 protein interacts with receptor-regulated Smads (R-Smads) Smads1-5, the co-Smad Smad4 and the inhibitory Smads (I-Smad6 and I-Smad7) in murine mesenchymal progenitors (66). TAK1 also affects subcellular distribution of Smad proteins and interferes with transactivation of R-Smads in reporter gene assays (66). We have previously reported that targeted inducible inactivation of TAK1 in skeletal muscle disrupts both Smad2/3 and Smad1/5/8 signaling (30). Interestingly, our experiments in the present study demonstrate that the phosphorylation of Smad1/5/8 as well as Smad2 is significantly increased in skeletal muscle expressing high levels of TAK1 activity. Furthermore, we found that the gene expression of various ligands and receptors of TGFβ/Myostatin/Activin A and BMP family of proteins was deregulated in skeletal muscle with hyperactivation of TAK1 (**Figure 8**). Even though the physiological significance and the mechanisms by which TAK1 regulates Smad1/5/8 and Smad2/3 signaling remain unknown, deregulation of Smad signaling could be another mechanism for the loss of muscle mass in response to high levels of TAK1 activation.

Our study also has some limitations. For example, we have investigated the effects of low and high levels of TAK1 activity in skeletal muscle at only one time point. It would be informative to understand the effects of forced activation of TAK1 at multiple time points after co-injection of TAK and TAB1 AAVs in skeletal muscle of mice. Moreover, we used AAV6 vectors that can also transduce non-muscle cells in skeletal muscle. There are also reports suggesting that AAV6 serotype can cause inflammation in the absence of transgene. However, we did not find any sign of inflammation in contralateral muscle transduced with equal amount of recombinant AAV6-GFP suggesting that it is the increased activation of TAK1, not the vector itself, which causes muscle injury and inflammation. Still use of AAV serotypes, such as MyoAAVs, which show higher tropism for skeletal muscle *in vivo* (67), may be more appropriate to understand the effect of forced activation of TAK1 on skeletal muscle. Since there is an increase in cellular infiltrate in skeletal muscle expressing high levels of TAK1/TAB1, the surge in gene expression and phosphorylation of some of the molecules could be attributed to the increased infiltration of non-muscle cells in muscle tissues. More investigations are needed to determine how hyperactivation of TAK1 affects the activation of various signaling cascades and gene expression of proinflammatory molecules in skeletal muscle ex vivo or in cultured muscle cells.

In summary, our study demonstrates that while low levels of TAK1 activation promote muscle growth, its excessive activation shifts the phenotype from beneficial to pathological, resulting in degenerative myopathy. These results suggest that inhibiting TAK1 activity could be an effective strategy to mitigate pro-inflammatory, fibrotic, and catabolic processes, particularly in conditions such as inflammatory myopathies and other neuromuscular disorders where TAK1 activation is elevated.

## Methods

### Sex as a biological variable

All experiments involving animals were conducted using male mice at 11-14 weeks of age. We do not know if there are any gender-related differences about the effects of forced activation of TAK1 on skeletal muscle.

### Animals

C57BL/6 mice were purchased from Jackson Labs. The mice were used at the age of 10-14 weeks. For intramuscular expression of TAK1 and TAB1, the mice were anaesthetized with isoflurane and AAV vector genomes ranging from 10^9^ to 10^11^ (in 30µl PBS) were injected in the TA and GA muscle of mice. All the animals were handled according to approved institutional animal care and use committee (IACUC) protocols (protocol # PR201900043) of the University of Houston.

### AAV vectors

AAVs (serotype 6) were custom generated by Vector Biolabs (Malvern, PA, USA). The AAVs expressed either GFP, Tak1 (ref seq #BC006665) or Tab1 (ref seq# BC054369) genes under a ubiquitous CMV promoter.

### Histology and morphometric analysis

TA muscle injected with AAV6-GFP or a combination of AAV6-TAK1 and AAV6-TAB1 was isolated from mice and sectioned using a microtome cryostat. To access muscle architecture and quantify myofiber cross-sectional area (CSA), 8-μm-thick transverse sections of TA muscle were stained with Hematoxylin and Eosin (H&E). Sirius red (StatLab) staining was performed on TA muscle section to visualize fibrosis. All stained and immunofluorescence labeled sections were examined, and images were captured using an inverted microscope (Nikon Eclipse Ti-2E Inverted Microscope), a digital camera (Digital Sight DS-Fi3, Nikon) and NIS Elements AR software (Nikon) at room temperature. Quantitative analysis was performed using NIH ImageJ software, and image levels were equally adjusted using Photoshop CS6 software (Adobe).

### Immunohistochemistry

Frozen TA muscle sections were fixed with 4% paraformaldehyde prepared in phosphate buffered saline (PBS), blocked in 2% bovine serum albumin in PBS for 1 h, followed by incubation with anti-dystrophin, anti-eMyHC, anti-Pax7, and anti-laminin in blocking solution at 4 °C overnight under humidified conditions. The sections were washed with PBS followed incubation with donkey anti-rabbit Alexa Fluor 555, goat anti-mouse Alexa Fluor 594, or 568 and goat anti-rabbit Alexa Fluor 488 secondary antibody for 1 h at room temperature. Finally, the sections were washed three times for 15 min with PBS. Nuclei were counterstained with DAPI. The slides were mounted using fluorescence medium (Vector Laboratories).

### Western Blot

GA muscle tissues expressing GFP or TAK1/TAB1 were isolated from mice and rinsed in PBS and homogenized in lysis buffer (50 mM Tris-Cl (pH 8.0), 200□mM NaCl, 50 mM NaF, 1mM dithiothreitol, 1 mM sodium orthovanadate, 0.3% IGEPAL, and protease inhibitors).

Approximately, 100□μg of total protein was resolved on each lane on 8-12% SDS-PAGE gel, transferred onto a nitrocellulose membrane, and probed using specific primary antibody (Supplemental **Table 1**). Bound antibodies were detected by secondary antibodies conjugated to horseradish peroxidase (Cell Signaling Technology). Signal detection was performed by an enhanced chemiluminescence detection reagent (Bio-Rad). Approximate molecular masses were determined by comparison with the migration of prestained protein standards (Bio-Rad).

### RNA isolation and qRT-PCR

Total RNA isolation and qRT-PCR were performed as previously described (68). Briefly, total RNA was extracted from GA muscle of mice using TRIzol reagent (Thermo Fisher Scientific) and RNeasy Mini Kit (Qiagen, Valencia, CA, USA) according to the manufacturers’ protocols. First-strand cDNA for PCR analyses was synthesized using a commercially available iScript cDNA Synthesis Kit (Bio-Rad Laboratories). The quantification of mRNA expression was performed using the SYBR Green dye (Bio-Rad SsoAdvanced - Universal SYBR Green Supermix) method on a sequence detection system (CFX384 Touch Real-Time PCR Detection System - Bio-Rad Laboratories). The sequence of the primers is described in Supplemental **Table 2**. Data normalization was accomplished with the endogenous control (β-actin), and the normalized values were subjected to a 2^−ΔΔCt^ formula to calculate the fold change between control and experimental groups.

### Statistical analyses and experimental design

The sample size was calculated using power analysis methods based on the standart deviation (s.d.) and effect size previously obtained from the experimental procedures employed in the study. A minimal of eight animals per group was calculated. Considering a likely drop-off effect of 10%, we set sample size of each group of six mice. For some experiments, five animals were sufficient to obtain Statistically significant differences. Animals with same sex and same age were employed to minimize physiological variability and to reduce s.d. from mean. The exclusion criteria for animals were established in consultation with a veterinarian and experimental outcomes. In case of death, skin injury, ulceration, sickness, or weight loss of > 10%, the animal was excluded from analysis. Tissue samples were excluded in cases such as freeze artifacts on histological sections or failure in extraction of RNA or protein of suitable quality and quantity. We included animals from different breeding cages by random allocation to the different experimental groups. All animal experiments were conducted in a blinded manner, using number codes until the final data analyses were performed. Statistical tests were used as described in the figure legends. Results are expressed as mean ± SEM. Statistical analyses used two-tailed Student’s t-test. A value of p ≤ 0.05 was considered statistically significant unless otherwise specified.

### Study approval

All animal procedures were conducted in strict accordance with the institutional guidelines and were approved by the Institutional Animal Care and Use Committee and Institutional Biosafety Committee of the University of Houston (PROTO201900043).

## Acknowledgements

This work was supported by the National Institute of Health grant AR081487 and CA294365 to AK.

## Authors’ contribution

A.K. designed the work. M.T.S. and A.R. performed all the experiments. M.T.S. wrote the first draft of the manuscript and A.R. and A.K. edited and finalized the manuscript.

## Notes

### Competing Interest Statement

The authors have declared no competing interest.

### Summary of Updates

There were some typos in the manuscript that have been fixed. The discussion section of the manuscript is improved. Figures 5 and 6 and supplemental Figure S4 have been revised to include additional blots.

## References

1. Larsson, L., Degens, H., Li, M., Salviati, L., Lee, Y. I., Thompson, W., Kirkland, J. L., and Sandri, M. (2019) Sarcopenia: Aging-Related Loss of Muscle Mass and Function. Physiol Rev 99, 427–511

2. Sandri, M. (2008) Signaling in muscle atrophy and hypertrophy. Physiology 23, 160–170

3. Bonaldo, P., and Sandri, M. (2013) Cellular and molecular mechanisms of muscle atrophy. Dis Model Mech 6, 25–39

4. Shin, J., Tajrishi, M. M., Ogura, Y., and Kumar, A. (2013) Wasting mechanisms in muscular dystrophy. Int J Biochem Cell Biol 45, 2266–2279

5. Blake, D. J., Weir, A., Newey, S. E., and Davies, K. E. (2002) Function and genetics of dystrophin and dystrophin-related proteins in muscle. Physiological reviews 82, 291–329

6. Emery, A. E. (2002) The muscular dystrophies. Lancet 359, 687–695

7. Wischnewski, S., Rausch, H. W., Ikenaga, C., Leipe, J., Lloyd, T. E., and Schirmer, L. (2025) Emerging mechanisms and therapeutics in inflammatory muscle diseases. Trends Pharmacol Sci 46, 249–263

8. Mastaglia, F. L., Garlepp, M. J., Phillips, B. A., and Zilko, P. J. (2003) Inflammatory myopathies: clinical, diagnostic and therapeutic aspects. Muscle Nerve 27, 407–425

9. Sartori, R., Romanello, V., and Sandri, M. (2021) Mechanisms of muscle atrophy and hypertrophy: implications in health and disease. Nat Commun 12, 330

10. Egerman, M. A., and Glass, D. J. (2014) Signaling pathways controlling skeletal muscle mass. Critical reviews in biochemistry and molecular biology 49, 59–68

11. Li, H., Malhotra, S., and Kumar, A. (2008) Nuclear factor-kappa B signaling in skeletal muscle atrophy. J Mol Med (Berl) 86, 1113–1126

12. Doyle, A., Zhang, G., Abdel Fattah, E. A., Eissa, N. T., and Li, Y. P. (2011) Toll-like receptor 4 mediates lipopolysaccharide-induced muscle catabolism via coordinate activation of ubiquitin-proteasome and autophagy-lysosome pathways. FASEB J 25, 99–110

13. Agca, S., and Kir, S. (2024) The role of interleukin-6 family cytokines in cancer cachexia. FEBS J 291, 4009–4023

14. Bonetto, A., Aydogdu, T., Jin, X., Zhang, Z., Zhan, R., Puzis, L., Koniaris, L. G., and Zimmers, T. A. (2012) JAK/STAT3 pathway inhibition blocks skeletal muscle wasting downstream of IL-6 and in experimental cancer cachexia. Am J Physiol Endocrinol Metab 303, E410–421

15. Sartori, R., and Sandri, M. (2015) Bone and morphogenetic protein signalling and muscle mass. Curr Opin Clin Nutr Metab Care 18, 215–220

16. Sartori, R., Schirwis, E., Blaauw, B., Bortolanza, S., Zhao, J., Enzo, E., Stantzou, A., Mouisel, E., Toniolo, L., Ferry, A., Stricker, S., Goldberg, A. L., Dupont, S., Piccolo, S., Amthor, H., and Sandri, M. (2013) BMP signaling controls muscle mass. Nature genetics 45, 1309–1318

17. Chen, J. L., Walton, K. L., Hagg, A., Colgan, T. D., Johnson, K., Qian, H., Gregorevic, P., and Harrison, C. A. (2017) Specific targeting of TGF-beta family ligands demonstrates distinct roles in the regulation of muscle mass in health and disease. Proc Natl Acad Sci U S A 114, E5266–E5275

18. Sartori, R., Gregorevic, P., and Sandri, M. (2014) TGFbeta and BMP signaling in skeletal muscle: potential significance for muscle-related disease. Trends Endocrinol Metab 25, 464–471

19. Dasgupta, A., Gibbard, D. F., Schmitt, R. E., Arneson-Wissink, P. C., Ducharme, A. M., Bruinsma, E. S., Hawse, J. R., Jatoi, A., and Doles, J. D. (2023) A TGF-beta/KLF10 signaling axis regulates atrophy-associated genes to induce muscle wasting in pancreatic cancer. Proc Natl Acad Sci U S A 120, e2215095120

20. Winbanks, C. E., Chen, J. L., Qian, H., Liu, Y., Bernardo, B. C., Beyer, C., Watt, K. I., Thomson, R. E., Connor, T., Turner, B. J., McMullen, J. R., Larsson, L., McGee, S. L., Harrison, C. A., and Gregorevic, P. (2013) The bone morphogenetic protein axis is a positive regulator of skeletal muscle mass. J Cell Biol 203, 345–357

21. Winbanks, C. E., Murphy, K. T., Bernardo, B. C., Qian, H., Liu, Y., Sepulveda, P. V., Beyer, C., Hagg, A., Thomson, R. E., Chen, J. L., Walton, K. L., Loveland, K. L., McMullen, J. R., Rodgers, B. D., Harrison, C. A., Lynch, G. S., and Gregorevic, P. (2016) Smad7 gene delivery prevents muscle wasting associated with cancer cachexia in mice. Sci Transl Med 8, 348ra398

22. Xu, Y. R., and Lei, C. Q. (2020) TAK1-TABs Complex: A Central Signalosome in Inflammatory Responses. Front Immunol 11, 608976

23. Mihaly, S. R., Ninomiya-Tsuji, J., and Morioka, S. (2014) TAK1 control of cell death. Cell Death Differ 21, 1667–1676

24. Choi, M. E., Ding, Y., and Kim, S. I. (2012) TGF-beta signaling via TAK1 pathway: role in kidney fibrosis. Semin Nephrol 32, 244–252

25. Ajibade, A. A., Wang, H. Y., and Wang, R. F. (2013) Cell type-specific function of TAK1 in innate immune signaling. Trends in immunology 34, 307–316

26. Singhirunnusorn, P., Suzuki, S., Kawasaki, N., Saiki, I., and Sakurai, H. (2005) Critical roles of threonine 187 phosphorylation in cellular stress-induced rapid and transient activation of transforming growth factor-beta-activated kinase 1 (TAK1) in a signaling complex containing TAK1-binding protein TAB1 and TAB2. J Biol Chem 280, 7359–7368

27. Yu, Y., Ge, N., Xie, M., Sun, W., Burlingame, S., Pass, A. K., Nuchtern, J. G., Zhang, D., Fu, S., Schneider, M. D., Fan, J., and Yang, J. (2008) Phosphorylation of Thr-178 and Thr-184 in the TAK1 T-loop is required for interleukin (IL)-1-mediated optimal NFkappaB and AP-1 activation as well as IL-6 gene expression. J Biol Chem 283, 24497–24505

28. Sato, S., Sanjo, H., Takeda, K., Ninomiya-Tsuji, J., Yamamoto, M., Kawai, T., Matsumoto, K., Takeuchi, O., and Akira, S. (2005) Essential function for the kinase TAK1 in innate and adaptive immune responses. Nat Immunol 6, 1087–1095

29. Hindi, S. M., Sato, S., Xiong, G., Bohnert, K. R., Gibb, A. A., Gallot, Y. S., McMillan, J. D., Hill, B. G., Uchida, S., and Kumar, A. (2018) TAK1 regulates skeletal muscle mass and mitochondrial function. JCI Insight 3, e98441

30. Roy, A., and Kumar, A. (2022) Supraphysiological activation of TAK1 promotes skeletal muscle growth and mitigates neurogenic atrophy. Nat Commun 13, 2201

31. Xu, D., Li, S., Wang, L., Jiang, J., Zhao, L., Huang, X., Sun, Z., Li, C., Sun, L., Li, X., Jiang, Z., and Zhang, L. (2021) TAK1 inhibition improves myoblast differentiation and alleviates fibrosis in a mouse model of Duchenne muscular dystrophy. J Cachexia Sarcopenia Muscle 12, 192–208

32. Xu, D., Zhao, L., Jiang, J., Li, S., Sun, Z., Huang, X., Li, C., Wang, T., Sun, L., Li, X., Jiang, Z., and Zhang, L. (2020) A potential therapeutic effect of catalpol in Duchenne muscular dystrophy revealed by binding with TAK1. J Cachexia Sarcopenia Muscle 11, 1306–1320

33. Kishimoto, K., Matsumoto, K., and Ninomiya-Tsuji, J. (2000) TAK1 mitogen-activated protein kinase kinase kinase is activated by autophosphorylation within its activation loop. J Biol Chem 275, 7359–7364

34. Srivastava, A., Mallela, K. M. G., Deorkar, N., and Brophy, G. (2021) Manufacturing Challenges and Rational Formulation Development for AAV Viral Vectors. J Pharm Sci 110, 2609–2624

35. Zhao, L., Wang, Q., Zhou, B., Zhang, L., and Zhu, H. (2021) The Role of Immune Cells in the Pathogenesis of Idiopathic Inflammatory Myopathies. Aging Dis 12, 247–260

36. Relaix, F., and Zammit, P. S. (2012) Satellite cells are essential for skeletal muscle regeneration: the cell on the edge returns centre stage. Development 139, 2845–2856

37. Yin, H., Price, F., and Rudnicki, M. A. (2013) Satellite cells and the muscle stem cell niche. Physiological reviews 93, 23–67

38. Li, J., Liang, C., Zhang, Z. K., Pan, X., Peng, S., Lee, W. S., Lu, A., Lin, Z., Zhang, G., Leung, W. N., and Zhang, B. T. (2017) TAK1 inhibition attenuates both inflammation and fibrosis in experimental pneumoconiosis. Cell Discov 3, 17023

39. Wang, W., Gao, W., Zhu, Q., Alasbahi, A., Seki, E., and Yang, L. (2021) TAK1: A Molecular Link Between Liver Inflammation, Fibrosis, Steatosis, and Carcinogenesis. Front Cell Dev Biol 9, 734749

40. Bale, S., Verma, P., Yalavarthi, B., Scarneo, S. A., Hughes, P., Amin, M. A., Tsou, P. S., Khanna, D., Haystead, T. A., Bhattacharyya, S., and Varga, J. (2023) Pharmacological inhibition of TAK1 prevents and induces regression of experimental organ fibrosis. JCI Insight 8

41. Akhmetshina, A., Palumbo, K., Dees, C., Bergmann, C., Venalis, P., Zerr, P., Horn, A., Kireva, T., Beyer, C., Zwerina, J., Schneider, H., Sadowski, A., Riener, M. O., MacDougald, O. A., Distler, O., Schett, G., and Distler, J. H. (2012) Activation of canonical Wnt signalling is required for TGF-beta-mediated fibrosis. Nat Commun 3, 735

42. Biressi, S., Miyabara, E. H., Gopinath, S. D., Carlig, P. M., and Rando, T. A. (2014) A Wnt-TGFbeta2 axis induces a fibrogenic program in muscle stem cells from dystrophic mice. Sci Transl Med 6, 267ra176

43. Brack, A. S., Conboy, M. J., Roy, S., Lee, M., Kuo, C. J., Keller, C., and Rando, T. A. (2007) Increased Wnt signaling during aging alters muscle stem cell fate and increases fibrosis. Science 317, 807–810

44. Aman, Y., Schmauck-Medina, T., Hansen, M., Morimoto, R. I., Simon, A. K., Bjedov, I., Palikaras, K., Simonsen, A., Johansen, T., Tavernarakis, N., Rubinsztein, D. C., Partridge, L., Kroemer, G., Labbadia, J., and Fang, E. F. (2021) Autophagy in healthy aging and disease. Nat Aging 1, 634–650

45. Hall, D. T., Griss, T., Ma, J. F., Sanchez, B. J., Sadek, J., Tremblay, A. M. K., Mubaid, S., Omer, A., Ford, R. J., and Bedard, N. (2018) The AMPK agonist 5 aminoimidazole 4 carboxamide ribonucleotide (AICAR), but not metformin, prevents inflammation associated cachectic muscle wasting. EMBO molecular medicine 10, e8307

46. Barbeau, J. E., Stark, I. S., Summers, A. E., Gold, A. B., Dray, S. D., and Thomson, D. M. (2024) Small-molecule AMPK activation preserves anabolic signaling in a C2C12 myotube cachexia model. Physiology 39, 2450

47. Raun, S. H., Ali, M. S., Han, X., Henríquez Olguín, C., Pham, T. C. P., Meneses Valdés, R., Knudsen, J. R., Willemsen, A. C. H., Larsen, S., and Jensen, T. E. (2023) Adenosine monophosphate activated protein kinase is elevated in human cachectic muscle and prevents cancer induced metabolic dysfunction in mice. Journal of cachexia, sarcopenia and muscle 14, 1631–1647

48. Setiawan, T., Sari, I. N., Wijaya, Y. T., Julianto, N. M., Muhammad, J. A., Lee, H., Chae, J. H., and Kwon, H. Y. (2023) Cancer cachexia: molecular mechanisms and treatment strategies. Journal of hematology & oncology 16, 54

49. White, J. P., Baynes, J. W., Welle, S. L., Kostek, M. C., Matesic, L. E., Sato, S., and Carson, J. A. (2011) The regulation of skeletal muscle protein turnover during the progression of cancer cachexia in the ApcMin/+ mouse. PloS one 6, e24650

50. Joseph, G. A., Wang, S. X., Jacobs, C. E., Zhou, W., Kimble, G. C., Tse, H. W., Eash, J. K., Shavlakadze, T., and Glass, D. J. (2019) Partial Inhibition of mTORC1 in Aged Rats Counteracts the Decline in Muscle Mass and Reverses Molecular Signaling Associated with Sarcopenia. Mol Cell Biol 39

51. Tang, H., Inoki, K., Brooks, S. V., Okazawa, H., Lee, M., Wang, J., Kim, M., Kennedy, C. L., Macpherson, P. C. D., Ji, X., Van Roekel, S., Fraga, D. A., Wang, K., Zhu, J., Wang, Y., Sharp, Z. D., Miller, R. A., Rando, T. A., Goldman, D., Guan, K. L., and Shrager, J. B. (2019) mTORC1 underlies age-related muscle fiber damage and loss by inducing oxidative stress and catabolism. Aging Cell 18, e12943

52. Roy, A., Narkar, V. A., and Kumar, A. (2023) Emerging role of TAK1 in the regulation of skeletal muscle mass. Bioessays 45, e2300003

53. Ogura, Y., Hindi, S. M., Sato, S., Xiong, G., Akira, S., and Kumar, A. (2015) TAK1 modulates satellite stem cell homeostasis and skeletal muscle repair. Nat Commun 6, 10123

54. Roy, A., Koike, T. E., Joshi, A. S., Tomaz da Silva, M., Mathukumalli, K., Wu, M., and Kumar, A. (2023) Targeted regulation of TAK1 counteracts dystrophinopathy in a DMD mouse model. JCI Insight 8

55. Sciorati, C., Rigamonti, E., Manfredi, A. A., and Rovere-Querini, P. (2016) Cell death, clearance and immunity in the skeletal muscle. Cell Death Differ 23, 927–937

56. Hayden, M. S., and Ghosh, S. (2004) Signaling to NF-kappaB. Genes Dev 18, 2195–2224

57. Jackman, R. W., Cornwell, E. W., Wu, C. L., and Kandarian, S. C. (2013) Nuclear factor-kappaB signalling and transcriptional regulation in skeletal muscle atrophy. Exp Physiol 98, 19–24

58. Bilgic, S. N., Domaniku, A., Toledo, B., Agca, S., Weber, B. Z. C., Arabaci, D. H., Ozornek, Z., Lause, P., Thissen, J. P., Loumaye, A., and Kir, S. (2023) EDA2R-NIK signalling promotes muscle atrophy linked to cancer cachexia. Nature 617, 827–834

59. Karin, M. (2006) Role for IKK2 in muscle: waste not, want not. J Clin Invest 116, 2866–2868

60. Cai, D., Frantz, J. D., Tawa, N. E., Jr., Melendez, P. A., Oh, B. C., Lidov, H. G., Hasselgren, P. O., Frontera, W. R., Lee, J., Glass, D. J., and Shoelson, S. E. (2004) IKKbeta/NF-kappaB activation causes severe muscle wasting in mice. Cell 119, 285–298

61. Zhang, G., Jin, B., and Li, Y. P. (2011) C/EBPbeta mediates tumour-induced ubiquitin ligase atrogin1/MAFbx upregulation and muscle wasting. EMBO J 30, 4323–4335

62. Baracos, V. E., Martin, L., Korc, M., Guttridge, D. C., and Fearon, K. C. H. (2018) Cancer-associated cachexia. Nat Rev Dis Primers 4, 17105

63. Tang, H., Inoki, K., Lee, M., Wright, E., Khuong, A., Khuong, A., Sugiarto, S., Garner, M., Paik, J., DePinho, R. A., Goldman, D., Guan, K. L., and Shrager, J. B. (2014) mTORC1 promotes denervation-induced muscle atrophy through a mechanism involving the activation of FoxO and E3 ubiquitin ligases. Sci Signal 7, ra18

64. Castets, P., Lin, S., Rion, N., Di Fulvio, S., Romanino, K., Guridi, M., Frank, S., Tintignac, L. A., Sinnreich, M., and Ruegg, M. A. (2013) Sustained activation of mTORC1 in skeletal muscle inhibits constitutive and starvation-induced autophagy and causes a severe, late-onset myopathy. Cell Metab 17, 731–744

65. Castets, P., Rion, N., Theodore, M., Falcetta, D., Lin, S., Reischl, M., Wild, F., Guerard, L., Eickhorst, C., Brockhoff, M., Guridi, M., Ibebunjo, C., Cruz, J., Sinnreich, M., Rudolf, R., Glass, D. J., and Ruegg, M. A. (2019) mTORC1 and PKB/Akt control the muscle response to denervation by regulating autophagy and HDAC4. Nat Commun 10, 3187

66. Hoffmann, A., Preobrazhenska, O., Wodarczyk, C., Medler, Y., Winkel, A., Shahab, S., Huylebroeck, D., Gross, G., and Verschueren, K. (2005) Transforming growth factor-beta-activated kinase-1 (TAK1), a MAP3K, interacts with Smad proteins and interferes with osteogenesis in murine mesenchymal progenitors. J Biol Chem 280, 27271–27283

67. Tabebordbar, M., Lagerborg, K. A., Stanton, A., King, E. M., Ye, S., Tellez, L., Krunnfusz, A., Tavakoli, S., Widrick, J. J., Messemer, K. A., Troiano, E. C., Moghadaszadeh, B., Peacker, B. L., Leacock, K. A., Horwitz, N., Beggs, A. H., Wagers, A. J., and Sabeti, P. C. (2021) Directed evolution of a family of AAV capsid variants enabling potent muscle-directed gene delivery across species. Cell 184, 4919–4938 e4922

68. Hindi, S. M., Shin, J., Gallot, Y. S., Straughn, A. R., Simionescu-Bankston, A., Hindi, L., Xiong, G., Friedland, R. P., and Kumar, A. (2017) MyD88 promotes myoblast fusion in a cell-autonomous manner. Nat Commun 8, 1624

