## Supplemental Figures S1-4, Tables S1 and S2 for "Hyperactivation of TAK1 causes skeletal muscle pathology reminiscent of inflammatory myopathies"

This file contains Supplemental **Figures S1-S5** and **Tables S1** and **S2**

**FIGURE S1**

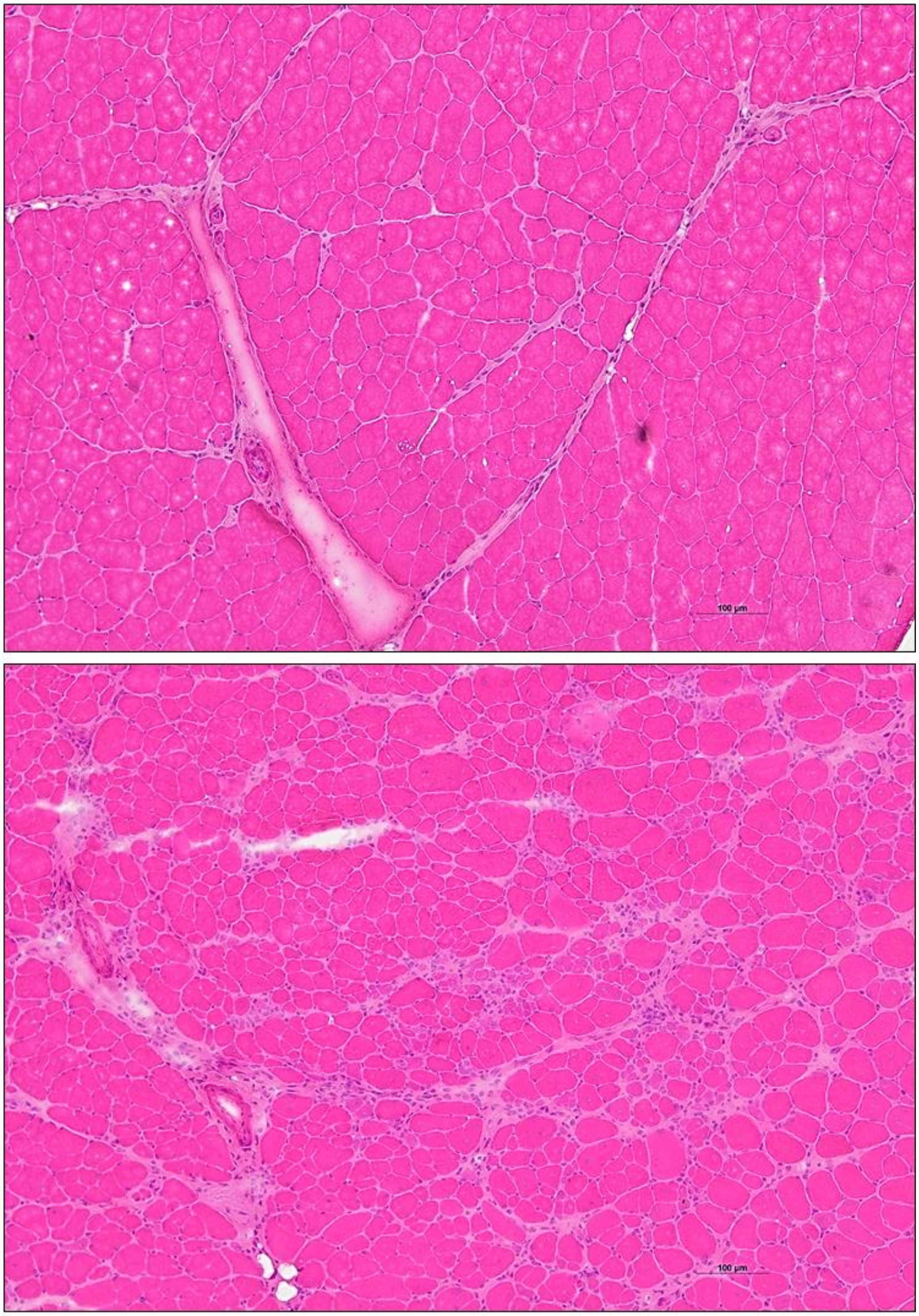


GFP (High)

TAK1/TAB1 (High)

H&E staining

**FIGURE S1. Hyperactivation of TAK1 causes myopathy in mice.** Uncropped 10X images of H&E-stained transverse sections of TA muscle of WT mice expressing high levels of GFP or TAK1/TAB1 protein. Scale bar: 100 µm, respectively.

**FIGURE S2**

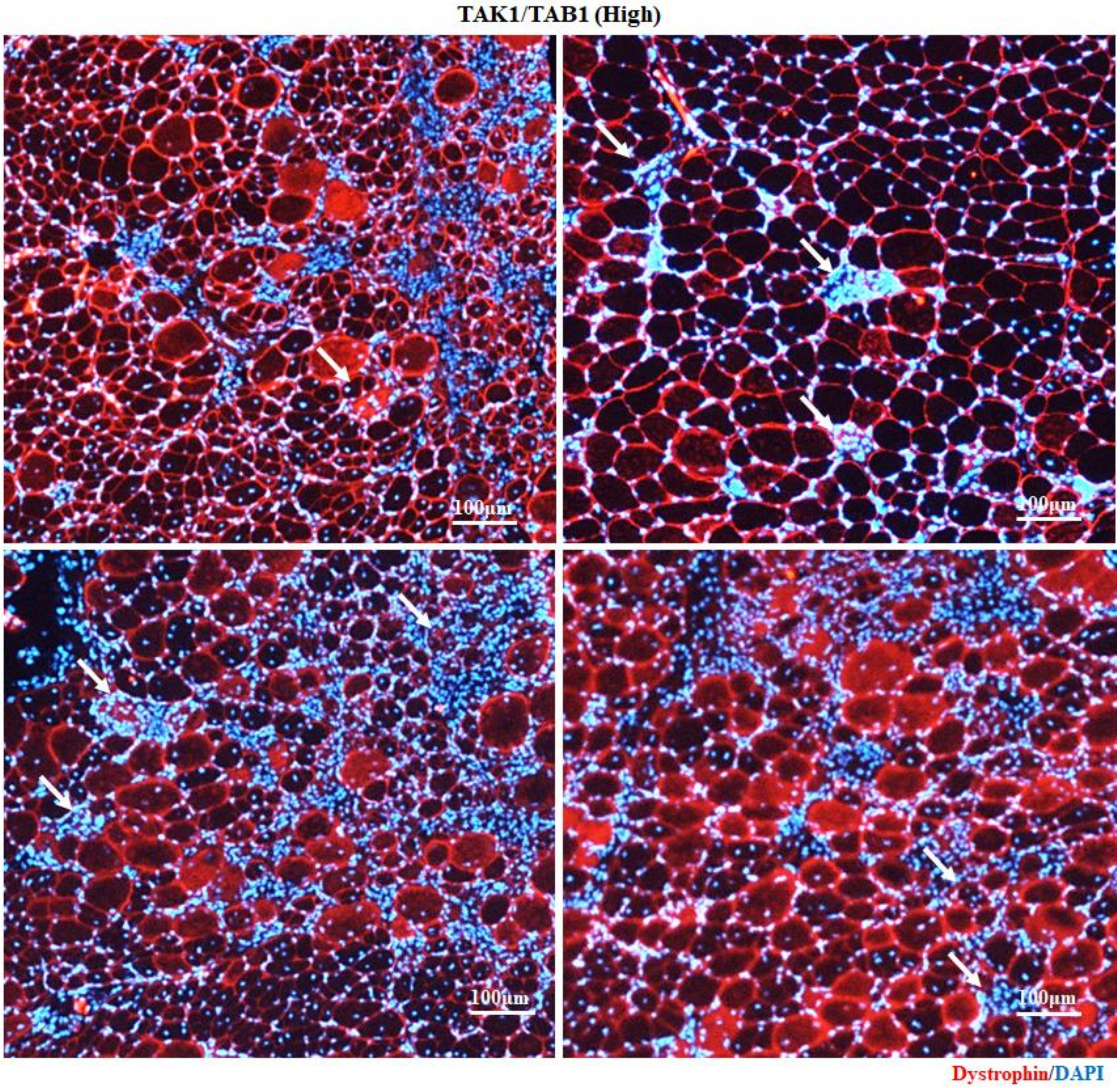


**FIGURE S2. Hyperactivation of TAK1 causes myofiber necrosis and inflammation in skeletal muscle.** Representative high magnification photomicrographs of transverse sections of TA muscle of mice expressing high levels of TAK1/TAB1 after immunostaining for dystrophin protein and DAPI staining. Scale bar: 100 µm. White arrows point to the necrotic myofibers filled with cellular infiltrate.

### FIGURE S3

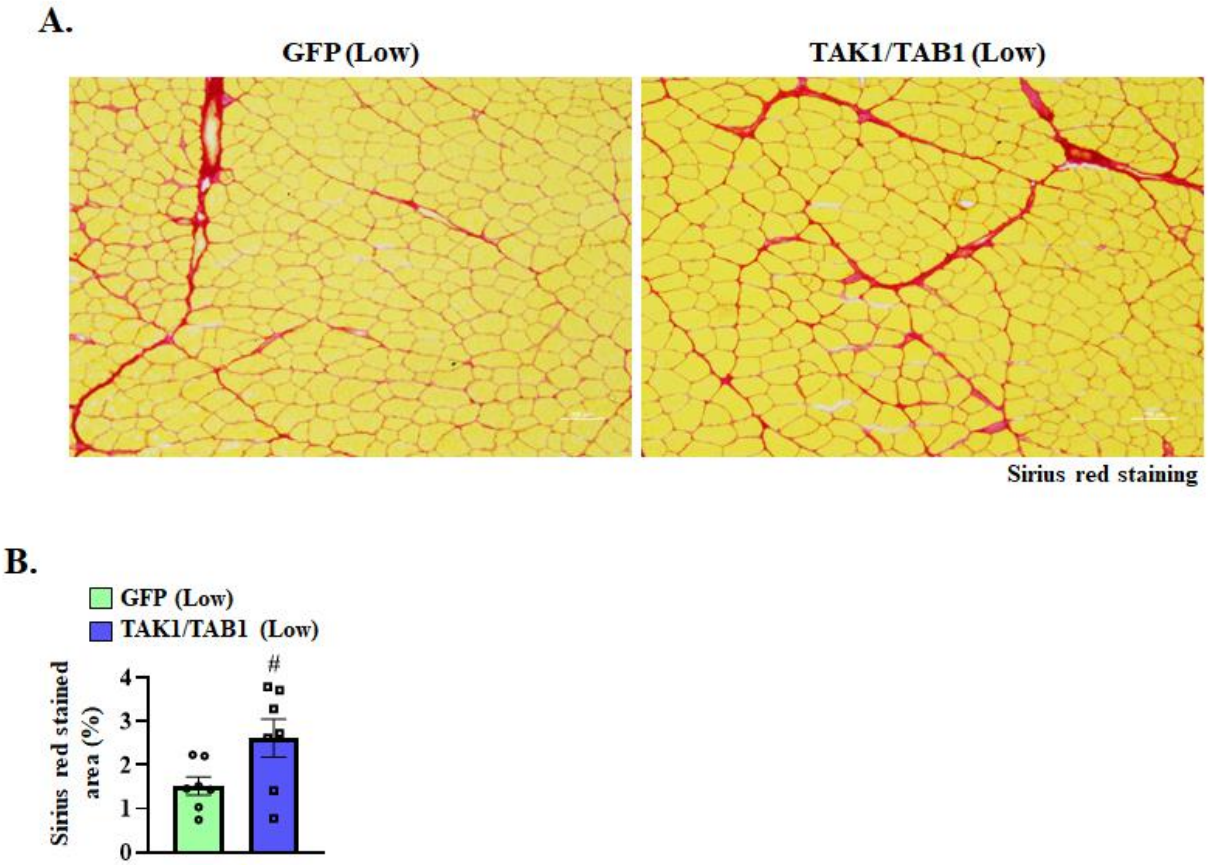


**FIGURE S3. Effects of low levels of TAK1 activity on fibrosis in skeletal muscle. (A)** Representative photomicrographs of Sirius red-stained TA muscle sections of WT mice expressing low levels of GFP or TAK1 and TAB1 protein. Scale bar: 100 µm. **(C)** Quantification of Sirius red-stained area of TA muscle sections expressing low levels of GFP or TAK1 and TAB1 protein. n= 6 mice in each group. All data are presented as mean ± SEM. #p ≤ 0.05, values significantly different from contralateral muscle expressing GFP analyzed by unpaired Student *t* test.

### FIGURE S4

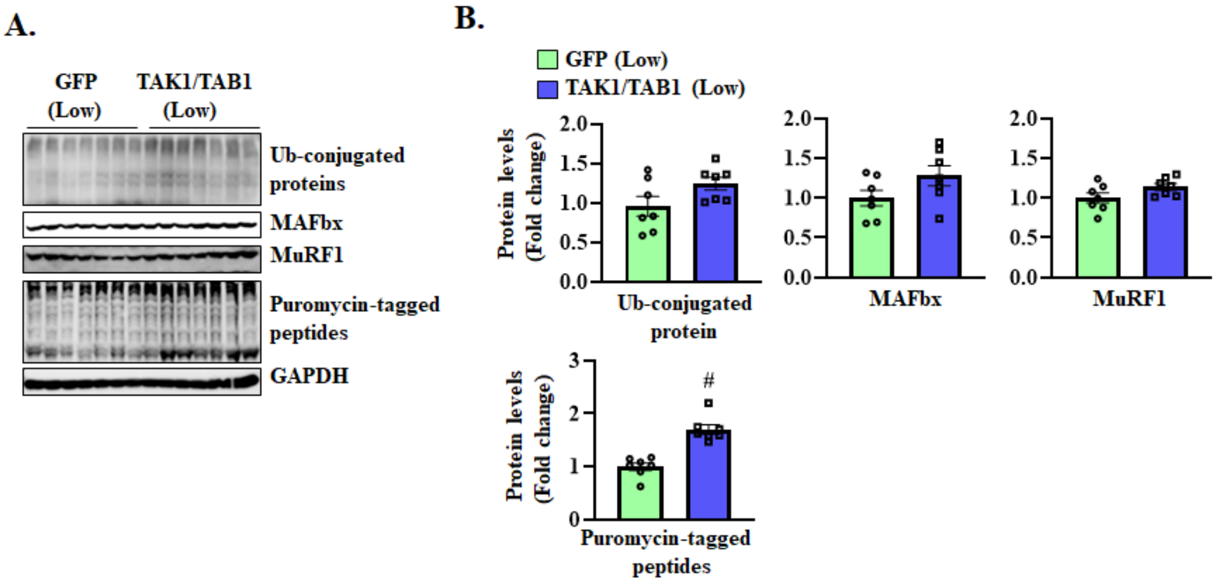


**FIGURE S4. Effect of low levels of TAK1 activation on ubiquitin proteasome system. (A)** Immunoblots and **(B)** densitometry analysis of protein levels of MAFbx and MuRF1, Ub-conjugated proteins, and puromycin-conjugated protein in GA muscle of mice expressing low levels of GFP or a combination of TAK1 and TAB1 protein. n= 7 mice in each group. All data are presented as mean ± SEM. #$p \leq 0.05$, values significantly different from contralateral muscle expressing GFP analyzed by unpaired Student *t* test. Western blots for A, and Figure 1C (in the main manuscript) were performed contemporaneously.

**FIGURE S5**

**1A.**

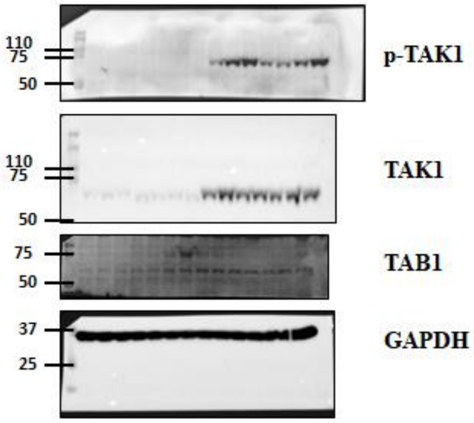


**1C.**

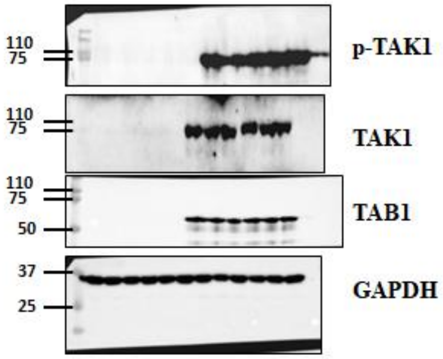


**3F.**

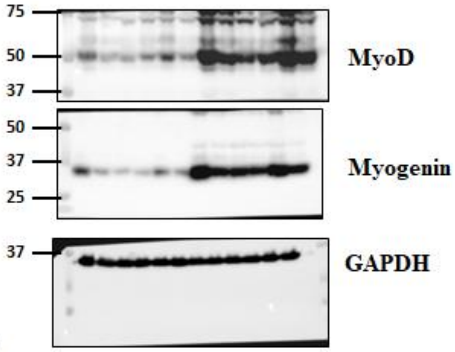


**4E.**

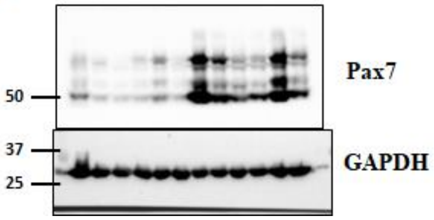


**5E.**

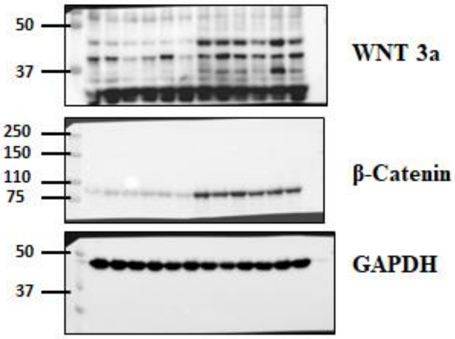


**6B.**

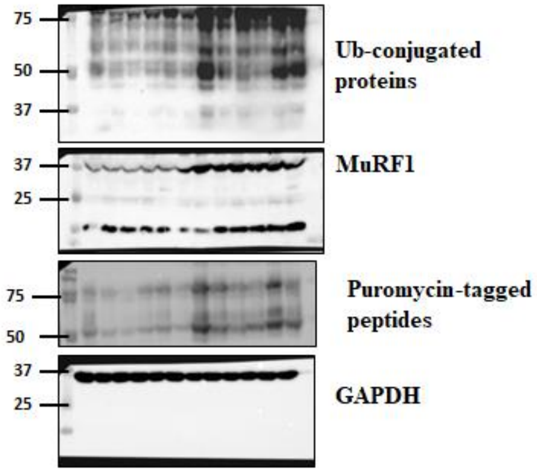


**6E.**

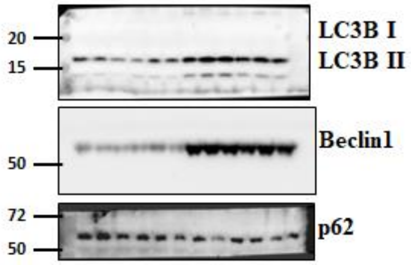

**FIGURE S5 (cont.)**

**7A.**

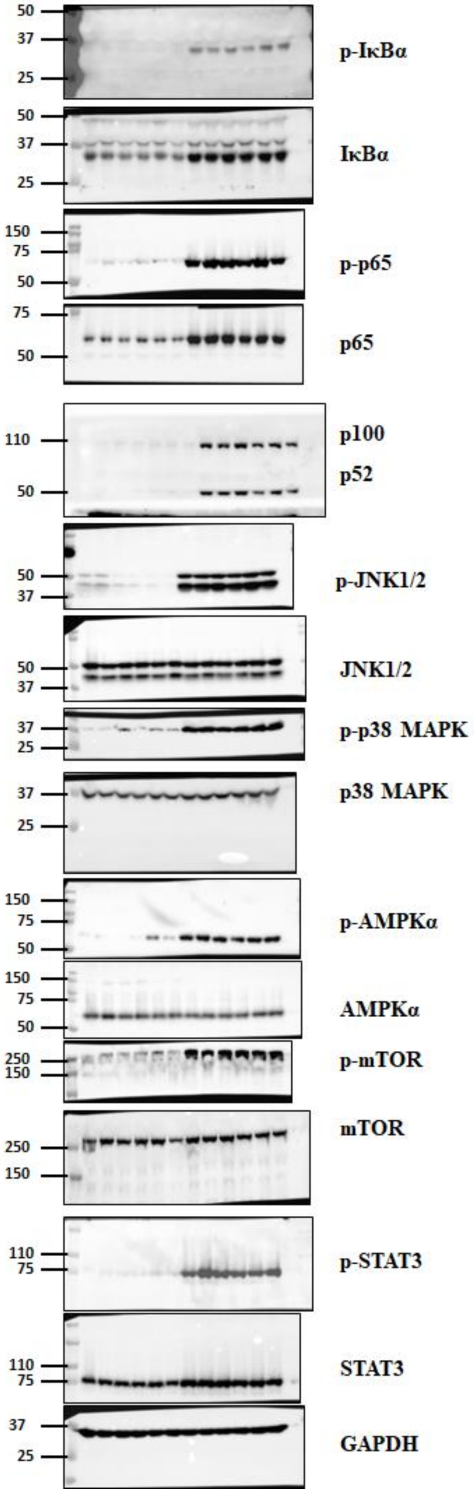


**8A.**

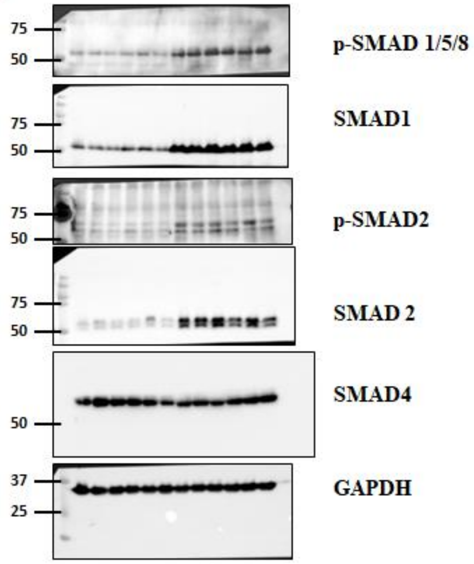


**S4A.**

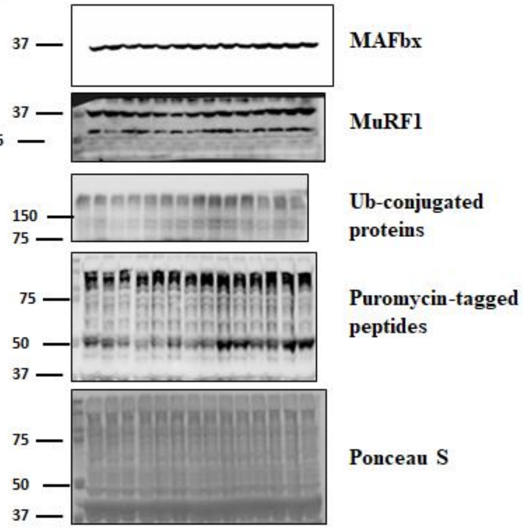


**FIGURE S5. Uncropped Western blot images.** The uncropped images of western blot are presented here

**Table S1.** List of antibodies used for Immunofluorescence and Western blot.

| Antibody | Source and Catalog no. |
|---|---|
| Monoclonal rabbit-anti-phospho-TAK1 | Invitrogen #MA5-15073 |
| Monoclonal rabbit-anti-total-TAK1 | Cell Signaling Technology, # 5206 |
| Monoclonal rabbit-anti-TAB1 | Cell Signaling Technology # 3226 |
| Monoclonal rabbit-anti-GAPDH | Cell Signaling Technology # 2118 |
| Polyclonal rabbit-anti-Dystrophin | Abcam, ab15277 |
| Monoclonal mouse-anti-eMyHC (embryonic) | DSHB # F1.652 |
| Monoclonal mouse-anti-MyoD | Santa Cruz Biotechnology # 377460 |
| Monoclonal mouse-anti-Myogenin | DSHB # F5D |
| Monoclonal mouse-anti-Pax7 | DSHB # PAX7 |
| Polyclonal rabbit-anti-Laminin | Sigma, L9393 |
| Polyclonal rabbit-anti-Wnt3a | Cell Signaling Technology, # 2391 |
| Monoclonal rabbit-anti-β-Catenin | Cell Signaling Technology, # 9582 |
| Donkey anti-Rabbit IgG Alexa Fluor 555 | Invitrogen # A31572 |
| Goat anti-Rabbit IgG Alexa Fluor 488 | Life Technologies # 11034 |
| Goat anti-Mouse IgG1 Alexa Fluor 568 | Life Technologies # A21124 |
| Goat anti-Mouse IgG2 Alexa Fluor 594 | Life Technologies # A211135 |
| Monoclonal mouse-anti-Ubiquitin | Santa Cruz Biotechnology, sc-8017 |
| Monoclonal mouse-anti-Puromycin | Millipore, MABE343 |
| Polyclonal goat-anti-MuRF1 | R&D Systems, AF5366 |
| Monoclonal rabbit-anti-LC3B | Cell Signaling Technology, # 3868 |
| Monoclonal rabbit-anti-Beclin-1 | Cell Signaling Technology, # 3495 |
| Polyclonal rabbit-anti-p62 | MBL #PM045 |
| Monoclonal rabbit-anti-p-IkBα | Cell Signaling Technology, # 2859 |
| Monoclonal rabbit-anti-IkBα | Cell Signaling Technology, # 4812 |
| Monoclonal rabbit-anti-phospho-p65 NF-κB | Cell Signaling Technology # 3033 |
| Monoclonal rabbit-anti-total-p65 NF-κB | Cell Signaling Technology # 8242 |
| Polyclonal rabbit-anti-NF-κB p100/p52 | Cell Signaling Technology # 4882 |
| Monoclonal rabbit-anti-p-JNK | Cell Signaling Technology # 4668 |
| Polyclonal rabbit-anti-JNK | Cell Signaling Technology # 9252 |
| Monoclonal rabbit-anti-phospho-p38 MAPK | Cell Signaling Technology, # 4511 |
| Polyclonal rabbit-anti-total-p38 MAPK | Cell Signaling Technology, # 9212 |
| Monoclonal rabbit-anti-phospho-AMPKα | Cell Signaling Technology # 2535 |
| Polyclonal rabbit-anti-total AMPKα | Cell Signaling Technology # 2532 |
| Monoclonal rabbit-anti-phospho-STAT3 (Y705) | Cell Signaling Technology # 9145 |
| Monoclonal rabbit-anti-STAT3 | Cell Signaling Technology # 30835 |
| Polyclonal rabbit-anti-phospho-mTOR | Cell Signaling Technology # 2971 |
| Polyclonal rabbit-anti-total-mTOR | Cell Signaling Technology # 2972 |
| Monoclonal rabbit-anti-phospho-Smad1/5/9 | Cell Signaling Technology # 13820 |
| Monoclonal rabbit-anti-total-Smad1 | Cell Signaling Technology # 6944 |
| Monoclonal rabbit-anti-phospho-Smad2 | Cell Signaling Technology # 3108 |
| Monoclonal rabbit-anti-total-Smad2 | Cell Signaling Technology # 5339 |
| Polyclonal rabbit-anti-Smad4 | Cell Signaling Technology # 9515 |
| Polyclonal rabbit-anti-MAFbx | ECM Biosciences, AP2041 |

**Supplemental Table 2.** List of primers used for QRT-PCR analysis.

| **Gene Name** | **Forward primer (5'-3')** | **Reverse primer (5'-3')** |
|---|---|---|
| *Myh3* | ACATCTCTATGCCACCTTCGCTAC | GGGTCTTGGTTTCGTTGGGTAT |
| *Myod1* | TGGGATATGGAGCTTCTATCGC | GGTGAGTCGAAACACGGATCAT |
| *Myog* | CAT CCA GTA CAT TGA GCG CCT A | GAG CAA ATG ATC TCC TGG GTT G |
| *Pax7* | CAGTGTGCCATCTACCCATGCTTA | GGTGCTTGGTTCAAATTGAGCC |
| *Tnfsf12* | GCTACGACCGCCAGATTGGG | GCCAGCACACCGTTCACCAG |
| *Tnfrsf12a* | AAGTGCATGGACTGCGCTTCTT | GGAAACTAGAAACCAGCGCCAA |
| *Tnfrsf1a* | AACCAGTTCCAACGCTACCTGA | AGAAAGAACCCTGCATGGCA |
| *Tnfrsf1b* | TAAGTGCCATCCCAAGGACACTCT | CCCAGTGATGTCACTCCAACAATC |
| *Adgre1* | CGTCAGGTACGGGATGAATATAAG | CTATGCCATCCACTTCCAAGAT |
| *Il6* | ATGGCAATTCTGATTGTATG | TGGCTTTGTCTTTCTTGTTA |
| *Il1b* | CTCCATGAGCTTTGTACAAGG | TGCTGATGTACCAGTTGGGG |
| *Tgfb1* | CTGAACCAAGGAGACGGAATAC | GGGCTGATCCCGTTGATTT |
| *Tgfb2* | GGCTTTCATTTGGCTTGAGATG | CTTCGGGTGAGACCACAAATAG |
| *Tgfb3* | GTACATCTGCTCTAGGGAATTGG | CCAGGCAGTGCAAGATATGA |
| *Wnt3a* | GCACCACCGTCAGCAACAG | GGGTGGCTTTGTCCAGAACA |
| *Wnt4* | CTGGAGAAGTGTGGCTGTGA | GGACGTCCACAAAGGACTGT |
| *Wnt5a* | GGCATCAAGGAATGCCAGTA | GTACGTGAAGGCCGTCTCTC |
| *Wnt7a* | TGAAGAGGACCCAGTGACAGG | GGCGTACTGGTGTGTGTTGT |
| *Wnt11* | GTAGGGCCTTCGCTGACAT | CGATGGTGTGACTGATGGTG |
| *Fzd1* | GCCGGCTGAGCTTGGAACTT | AACCAAAGCAGCAGCAGCAGC |
| *Fzd2* | CATCTCCATCCCGCTGTGCA | AGCACAGGAAGAAGCGCAGCTC |
| *Fzd4* | GGCTACAACGTGACCAAGATGCC | GCACATTGGCACATAAACCGAAC |
| *Fzd6* | GCGGCGTTTGCTTCGTT | CACAGAGGCAGAAGGACGAAGT |
| *Fbxo32* | GTCGCAGCCAAGAAGAGAAAGA | TGCTATCAGCTCCAACAGCCTT |
| *Trim63* | TACTGCATCTCCATGCTGGTG | TGGCGTAGAGGGTGTCAAACTT |
| *Fbxo30* | TCGTGGAATGGTAATCTTGC | CCTCCCGTTTCTCTATCACG |
| *Map1lc3b* | CTGGTGAATGGGCACAGCATG | CGTCCGCTGGTAACATCCCTT |
| *Becn1* | TGAAATCAATGCTGCCTGGG | CCAGAACAGTATAACGGCAACTCC |
| *Atg5* | ATCAGACCACGACGGAGCGG | GGCGACTGCGGAAGGACAGA |
| *Atg12* | ACAAAGAAATGGGCTGTGGAGC | GCAGTAATGCAGGACCAGTTTACC |
| *Mstn* | AATCCACCACGGTGCTAATG | TTAGTGCTGTGTGTGTGGAG |
| *Fst288* | AACGTTGGGAGAGAGGATGA | GACAGGTTGAAAGTGTGCCT |
| *Gdf3* | CTG GGC TTC TTG TTC TTG TTT G | ACA CAG GAG GTA GAG GAA AGA |
| *Gdf11* | GAA ACA TTG TGC GCC ACA | CCA CAT TCG ACA GGT CAA AGA |
| *Gdf15* | ATG CTG ACA GAA AGG GAG GTA A | GCA CTG ATG TCA AAC ACG TAC C |
| *Acvr1* | CCCAACTCTGAAACGGACAT | GTGGTGTTGCATGGGTAATG |
| *Bmpr1a* | GAA AGA CCT GAT TGA CCA GTC C | CCC ATC CAT ACT TCT CCA TAG C |
| *Acvr1b* | AAGCTGAGAGTTGGGAGAAG | GGGCTTTAGACTTGGTCTGT |
| *Acvr1c* | AACTTTGCCAACAGCTAGTC | GGCAGAGAAGAATGTACACC |

| | | |
|---|---|---|
| *Inhba* | AGA ACG GGT ATG TGG AGA TAG A | GAC TCG GCA AAG GTG ATG AT |
| *Bmpr1b* | GAATACCAGCTTCCCTATCACG | TCTGCCTGAGACACTCATCACT |
| *Bmpr2* | TTGGACTCATCTACTGGGAGGT | TGGACACAAGAACCTGCATATC |
| *Bmp4* | GACTTCGAGGCGACACTTCTAC | CAGATGTTCTTCGTGATGGAAA |
| *Bmp7* | AGCTTCGTCAACCTAGTGGAAC | CTGGAGCACCTGATAGACTGTG |
| *Gdf6* | AAGACTTACTCCATTGCCGAGA | TCGTCCAGTCCTCTGTCTACAA |
| *Actb* | CAGGCATTGCTGACAGGATG | TGCTGATCCACATCTGCTGG |
